# Loss of STIM1 and STIM2 in salivary glands disrupts ANO1 function but does not induce Sjogren’s disease

**DOI:** 10.1101/2024.01.08.574702

**Authors:** Ga-Yeon Son, Anna Zou, Amanda Wahl, Kai Ting Huang, Manikandan Vinu, Saruul Zorgit, Fang Zhou, Larry Wagner, Youssef Idaghdour, David I. Yule, Stefan Feske, Rodrigo S. Lacruz

**Affiliations:** Department of Molecular Pathobiology, New York University College of Dentistry, New York, USA; Department of Pharmacology and Physiology, University of Rochester, Rochester, New York, USA; Biology Program, Division of Science and Mathematics, New York University Abu Dhabi, Abu Dhabi, United Arab Emirates; Department of Pathology, New York University Grossman School of Medicine, New York, New York, USA

## Abstract

Sjogren’s disease (SjD) is an autoimmune disease characterized by xerostomia (dry mouth), lymphocytic infiltration into salivary glands and the presence of SSA and SSB autoantibodies. Xerostomia is caused by hypofunction of the salivary glands and has been involved in the development of SjD. Saliva production is regulated by parasympathetic input into the glands initiating intracellular Ca^2+^ signals that activate the store operated Ca^2+^ entry (SOCE) pathway eliciting sustained Ca^2+^ influx. SOCE is mediated by the STIM1 and STIM2 proteins and the ORAI1 Ca^2+^ channel. However, there are no studies on the effects of lack of STIM1/2 function in salivary acini in animal models and its impact on SjD. Here we report that male and female mice lacking *Stim1* and *Stim2* (*Stim1/2^K14Cre^*) in salivary glands showed reduced intracellular Ca^2+^ levels via SOCE in parotid acini and hyposalivate upon pilocarpine stimulation. Bulk RNASeq of the parotid glands of *Stim1/2^K14Cre^* mice showed a decrease in the expression of *Stim1/2* but no other Ca^2+^ associated genes mediating saliva fluid secretion. SOCE was however functionally required for the activation of the Ca^2+^ activated chloride channel ANO1. Despite hyposalivation, ageing *Stim1/2^K14Cre^* mice showed no evidence of lymphocytic infiltration in the glands or elevated levels of SSA or SSB autoantibodies in the serum, which may be linked to the downregulation of the toll-like receptor 8 (*Tlr8*). By contrast, salivary gland biopsies of SjD patients showed increased *STIM1* and *TLR8* expression, and induction of SOCE in a salivary gland cell line increased the expression of *TLR8*. Our data demonstrate that SOCE is an important activator of ANO1 function and saliva fluid secretion in salivary glands. They also provide a novel link between SOCE and TLR8 signaling which may explain why loss of SOCE does not result in SjD.

## Introduction

Sjogren’s disease (SjD) or Sjogren’s syndrome, is a chronic autoimmune disease characterized by the infiltration of T and B cells into salivary and lacrimal glands, and by loss of function of the glands resulting in sicca symptoms including dry mouth (xerostomia) (1, 2). SjD is more prevalent in females and occurs in ∼0.5% of the population (3–5). Disease diagnosis includes the evaluation of ocular and oral dryness, minor salivary gland biopsy and the detection of antibodies such as SS-A and SS-B (6–8). The vast majority of patients (90%) present with oral and or ocular dryness affecting quality of life, but extraglandular (systemic) symptoms may also be present (9, 10). The pathogenesis of SjD remains unclear and the cause of salivary fluid loss has not been identified (1, 7, 11). Factors associated with the disease include genetic predisposition and viral infections (8). A common feature of SjD is the slow and progressive course of the disease leading to cumulative tissue damage and mild symptoms before the diagnosis is made (12) suggesting that functional salivary gland defects may precede the development of SjD (13, 14).

Saliva performs important functions in the maintenance of oral health including the lubrication of oral mucosa, pH buffering and supporting enamel mineralization (15). Salivary acini are secretory epithelia that generate the fluid component of saliva in response to transcellular salt gradients regulated by intracellular Ca^2+^ signals (16, 17). All secretion is initiated by parasympathetic nervous input to the gland which activates the G-protein coupled muscarinic receptors (M1/M3) (18, 19) driving the release of Ca^2+^ pools from the endoplasmic reticulum (ER) via the inositol 1,4,5-trisphosphate receptor (IP3R) (20). A reduction in luminal ER Ca^2+^ activates store-operated Ca^2+^ entry (SOCE) to enable sustained Ca^2+^ influx into acinar cells (17). SOCE is mediated by Ca^2+^ release-activated Ca^2+^ (CRAC) channels that are formed by the plasma membrane channel ORAI1 and activated by the ER resident Ca^2+^ sensor proteins STIM1 and STIM2 (21). In salivary glands, the Ca^2+^ conducting transient receptor potential canonical channels (TRPC1) also contribute to Ca^2+^ influx (22), although ORAI1 is the primary channel activated by loss of ER Ca^2+^ (23). Global increases in cytosolic Ca^2+^ are important for the activation of ion channels as well as transporters that play a role in fluid secretion (16, 17, 24). In salivary acini, loss of function of the Ca^2+^-activated chloride channel ANO1 (encoded by *TMEM16A*) affects saliva fluid secretion (25), but if SOCE is necessary for ANO1 function in salivary glands is unknown.

Patients with mutations in *STIM1* and *ORAI1* present with a defect in eccrine sweat gland function (26). SOCE-deficient mice showed decreased sweat production because SOCE is necessary for the activation of ANO1 and chloride secretion (27). Further, we also reported that the outer appearance of the enamel of mice lacking *Stim1* and *Stim2* in epithelial ameloblast cells (*Stim1/2^K14Cre^*) was cracked, suggestive of a dry oral environment possibly because of limited saliva production (28). Despite the relevance of SOCE in fluid production in exocrine glands (11, 29), there are no reports on the role of *Stim1* and *Stim2* in vivo in mouse salivary glands or its impact on SjD. To address this question, and because loss of salivary gland function has been associated with the development of SjD, we investigated salivary gland function and SjD markers in *Stim1/2^K14Cre^* mice. We show that pilocarpine-stimulated saliva in male and female *Stim1/2^K14Cre^*mice was lower than in control mice and SOCE was significantly reduced. SOCE mediated Ca^2+^ influx and ANO1 currents in parotid glands were significantly decreased. However, *Stim1/2^K14Cre^*mice did not show lymphocytic infiltration or increased expression of SS autoantibodies. Our results identify an important role of STIM1/2 in saliva production via the activation of ANO1.

## Results

### *Stim1/2^K14Cre^* mice hyposalivate

To address if saliva production of *Stim1/2^K14Cre^* mice was affected, we stimulated saliva using the muscarinic receptor agonist pilocarpine in male and female *Stim1/2^K14Cre^* mice and wild type (WT) littermates. Male and female mice were weighted prior to collecting saliva on the same cohort of mice on a longitudinal study for over 22 weeks at intervals of approximately four weeks. Saliva volume was corrected by weight. We show that female and male *Stim1/2^K14Cre^* mice have decreased saliva production overall (Figure 1A and 1B) but do not have statistically lower body weight compared to controls (Figure 1C and 1D). The total volume of saliva in female *Stim1/2^K14Cre^* mice was significantly lower than in WT female mice starting at 17 weeks of age but in males this difference was significant at 30 weeks (Figure 1A and 1B). When corrected for body weight, these differences are maintained in females but become significantly different in males only at 34 weeks (Figure 1E and 1F). These data indicated that ageing *Stim1/2^K14Cre^* mice produced less saliva than WT mice.

**Figure 1.**
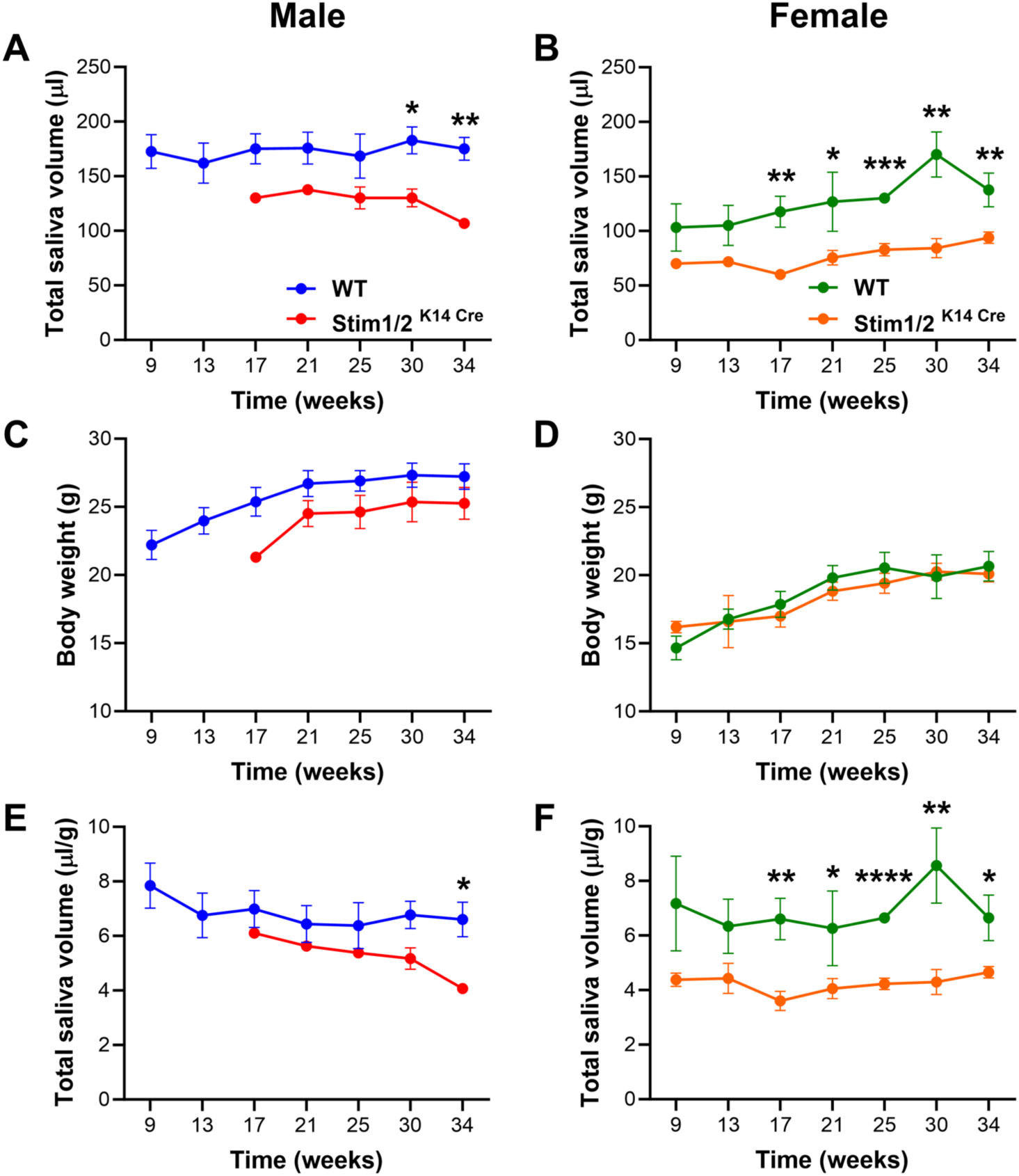
Mice lacking Stim1 and Stim2 in salivary glands hyposalivate: Male and female *Stim1/2^K14Cre^* mice and age-matched WT littermates were stimulated with pilocarpine to measure saliva fluid. The same animal cohorts were used throughout the longitudinal study and saliva was collected at each time point indicated. **(A-B)** Average of total saliva fluid collected per mouse at each time point in males (A) and females (B). **(C-D)** Body weigh average (in grams) in males (C) and females (D). N=at least 4 animals at each time point. **(E-F)** Average of saliva fluid corrected by body weight in males (E) and females (F). Data represent the mean ±SEM of values obtained from at least 4 animals per group at each time point. (\**P* < 0.5, \*\**P* < 0.01, \*\*\**P* < 0.001, \*\*\*\**P* < 0.0001, Two-way ANOVA followed by Tukey’s post-hoc test).

### SOCE is decreased in parotid acini of *Stim1/2^K14Cre^* mice

To address if Ca^2+^ signaling in parotid acinar cells was affected by the loss of *Stim1* and *Stim2*, we isolated parotid glands from 40-week-old male and female *Stim1/2^K14Cre^* mice and WT littermates. Dissociated salivary acini were stimulated with cyclopiazonic acid (CPA), a reversible inhibitor of the sarco-endoplasmic reticulum Ca^2+^-ATPase (SERCA) (30), to passively deplete the ER Ca^2+^ stores and stimulate SOCE by the readdition of a Ca^2+^ containing solution. Parotid acini of *Stim1/2^K14Cre^* mice showed a significant reduction in Ca^2+^ transients mediated by SOCE compared to WT littermates (Figure 2A and 2C). The peak of Ca^2+^ influx was significantly lower in parotid acini of both male and female *Stim1/2^K14Cre^* mice (Figure 2B and 2D). Males showed a reduction in ER Ca^2+^ release for unknown reasons (Figure 2B), absent in females (Figure 2D). These data highlight that loss of *Stim1* and *Stim2* results in a significant reduction in Ca^2+^ signaling via SOCE in parotid glands.

**Figure 2.**
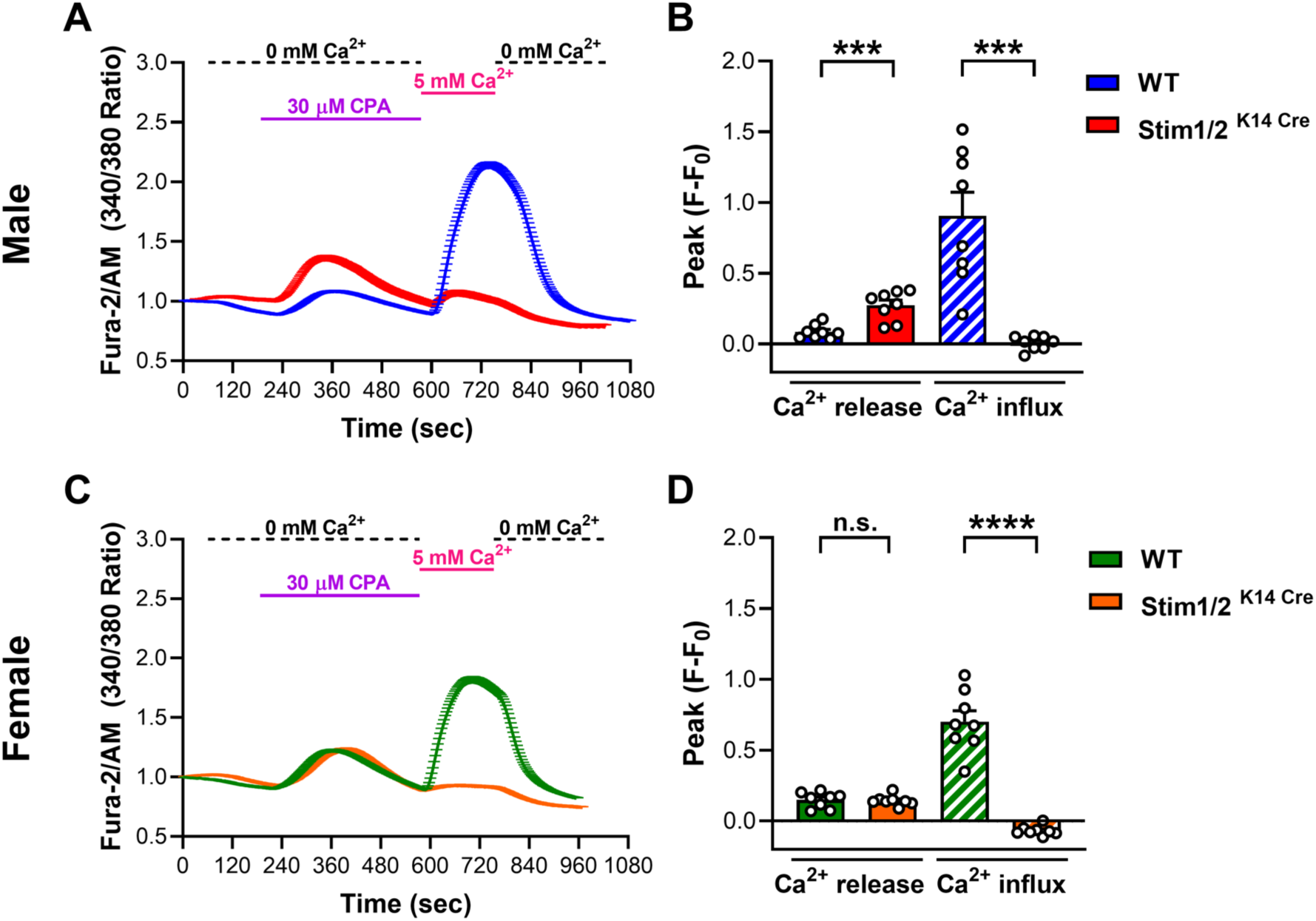
SOCE is impaired in parotid salivary glands of mice lacking Stim1 and Stim2. (A) Original Ca^2+^ traces of isolated parotid gland preparations of male *Stim1/2^K14Cre^* mice and age-matched WT littermates loaded with Fura 2-AM. Cells were treated with CPA (30 µM) to stimulate SOCE in Ca^2+^-free solutions before the readdition of 5 mM Ca^2+^. **(B)** Quantification of the ER Ca^2+^ release and SOCE peaks (F-F_o_). **(C and D)** Same as A and B but measured in parotid glands isolated from female *Stim1/2^K14Cre^* mice and age-matched WT littermates. Data in A-D represent the mean ±SEM from n = 3 different experiments. Each experiment sampled cells obtained from 1 WT mouse and 1 *Stim1/2^K14Cre^* mice littermate. Total cells: WT = 656, *Stim1/2^K14Cre^* mice = 777. (\*\*\**P* < 0.001, \*\*\*\**P* < 0.0001. n.s. = not significant. Two-tailed un-paired Student’s *t*-test).

### Genes involved in saliva production pathway are unaffected in *Stim1/2^K14Cre^* mice

To analyze the broader effects of SOCE deficiency in salivary glands, we isolated RNA from whole parotid glands of 40-week-old female *Stim1/2^K14Cre^* and WT mice and analyzed global gene expression by RNA sequencing (RNAseq). We used only female mice because hyposalivation was more prominent in females than in male (Figure1) and because SjD is more prevalent in females (3). A total of 140 genes were differentially expressed in *Stim1/2^K14Cre^* vs. WT mice (P < 0.05, FDR ≥ 1.2) with ∼70% being downregulated (Figure 3A and 3B, and Supplemental Figure 1). We specifically analyzed the expression of genes involved in the physiological control of saliva fluid production (11, 23). We found that the expression of *Stim1* and *Stim2* was significantly decreased, as expected (Figure 3C), whereas the expression of *Orai1* was unchanged. The levels of *Trpc1,* which has been reported to contribute to Ca^2+^ influx in salivary glands (22), was also unaffected (Figure 3C). Salivary glands express two main isoforms of the inositol 1,4,5-trisphosphate receptor, the type 2 (*Itp3r2)* and type 3 *(Itp3r3)* (31), that are essential in the release of intracellular Ca^2+^ (32), but their expression was not significantly affected in *Stim1/2^K14Cre^* mice, as well as other genes involved in Ca^2+^ signaling (Supplemental Figure 2). Similarly, the *Ano1* gene, encoding the Ca^2+^ activated chloride channel Anoctamin 1, was not differentially expressed (Figure 3C), and the water channel aquaporin 5 (*Aqp5*) channel involved in water transport in salivary glands showed a slight but non-significant increase in expression in *Stim1/2^K14Cre^* mice (Figure 3C). To further validate the RNASeq data, we performed quantitative reverse transcription polymerase chain reaction (qRT-PCR) analysis of the abovementioned genes, which confirmed the RNASeq results (Figure 3D). Western blot analysis of whole parotid glands showed lack of STIM1 expression in *Stim1/2^K14Cre^* mice whereas ANO1 and AQP5 levels were similar (Figure 3E-3H).

**Figure 3.**
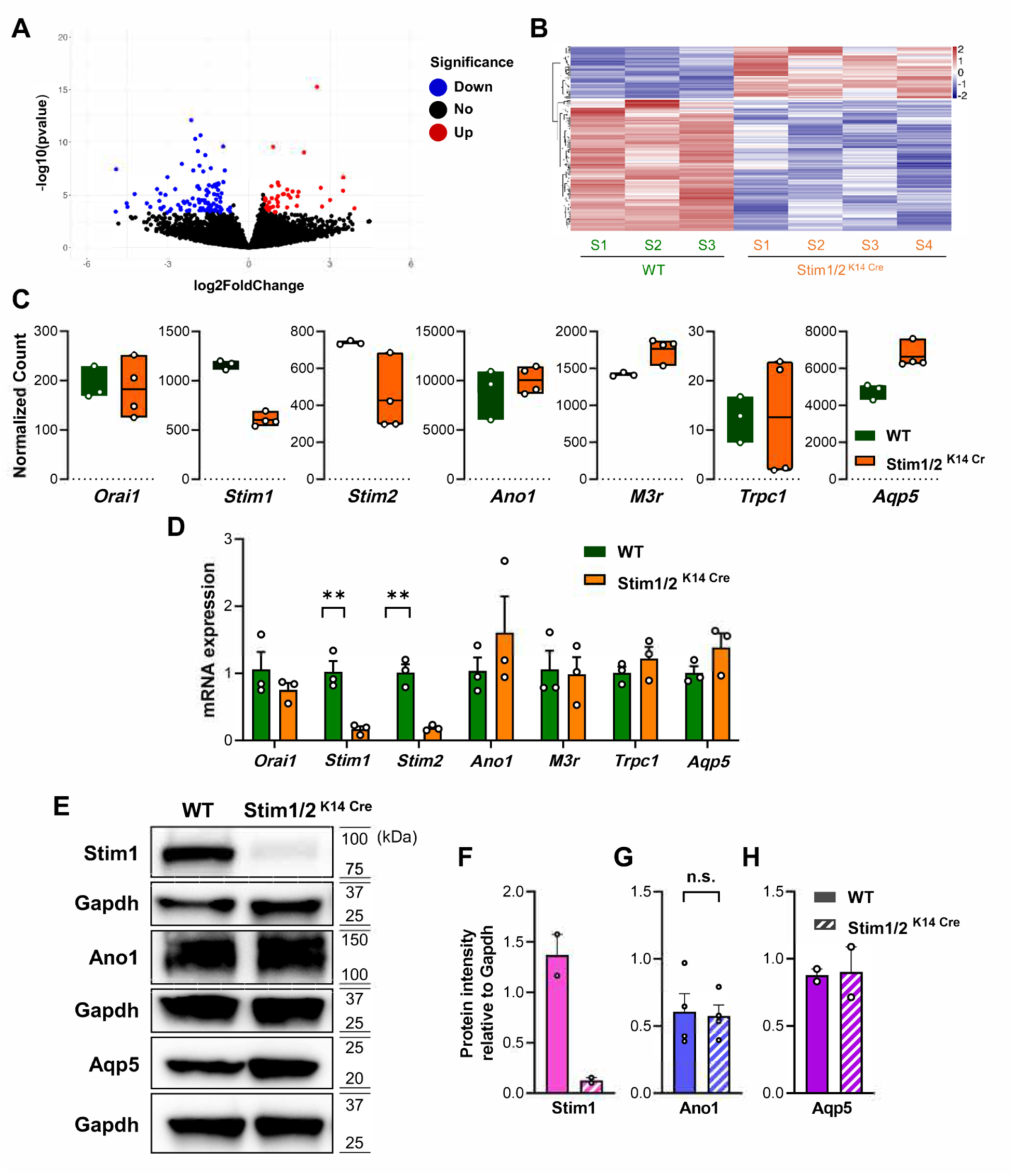
Ca^2+^ signaling genes are unaffected in parotid salivary glands of mice lacking Stim1 and Stim2. **(A)** Volcano plot of statistical significance (shown as the negative logarithm of the P value on the y axis) versus magnitude of differential gene expression (shown as the log base 2 of magnitude of mean expression difference on the x axis) between *Stim1/2^K14cre^* and WT mice. Differentially expressed genes (*P* < 0.05, One-way ANOVA) are shown in red (upregulation) and blue (downregulation). **(B)** Heatmap using two-way hierarchical clustering of WT and *Stim1/2^K14cre^* parotid glands of based on the criteria described in the text. **(C)** Normalized counts of genes involved in Ca^2+^ signaling in salivary glands such as *Orai1*, *Stim1*, *Stim2*, *Ano1*, *M3r*, *Trpc1*, and *Aqp5* between *Stim1/2^K14cre^* and WT mice. Normalized counts were generated by scaling raw count values to account for sequencing depth using DEseq2 (1.40.2). **(D)** qRT-PCR analysis of genes involved in Ca^2+^ signaling in salivary glands. Data represent the means (±SEM) of n = 3 mice per group (\*\**P* < 0.01, Two-way ANOVA followed by Tukey). **(E)**. Western blot analysis of protein expression levels in WT and *Stim1/2^K14cre^* parotid glands. **(F-H)** Densitometric analysis of protein expression levels based on two independent experiments analyzing STIM1 and AQP5, or three independent experiments for ANO1, all relative to GAPDH. Statistical significance was determined using a two-tailed Student’s t test. n.s. = not significant.

Because salivary fluid secretion in acinar cells relies on a transepithelial ion gradient from basal to apical poles (16), we addressed if the subcellular localization of proteins involved in ion transport were altered in parotid acini of *Stim1/2^K14Cre^* mice. We performed immunofluorescence staining and found that STIM1 expression is only observed in WT mice with strong cytoplasmic signals found in ductal cells and weaker signals in acinar cells (Figure 4A). Orai1 was localized to the lateral pole of acinar cells (Figure 4B). AQP5 and ANO1 are found at the apical pole of salivary acini of WT mice (33, 34) and in parotid acini of *Stim1/2^K14Cre^* mice this localization did not change (Figure 4C and 4D). These data suggest that loss of *Stim1* and *Stim2* in salivary glands does not evoke significant changes in the localization of proteins involved in the production of salivary fluid.

**Figure 4.**
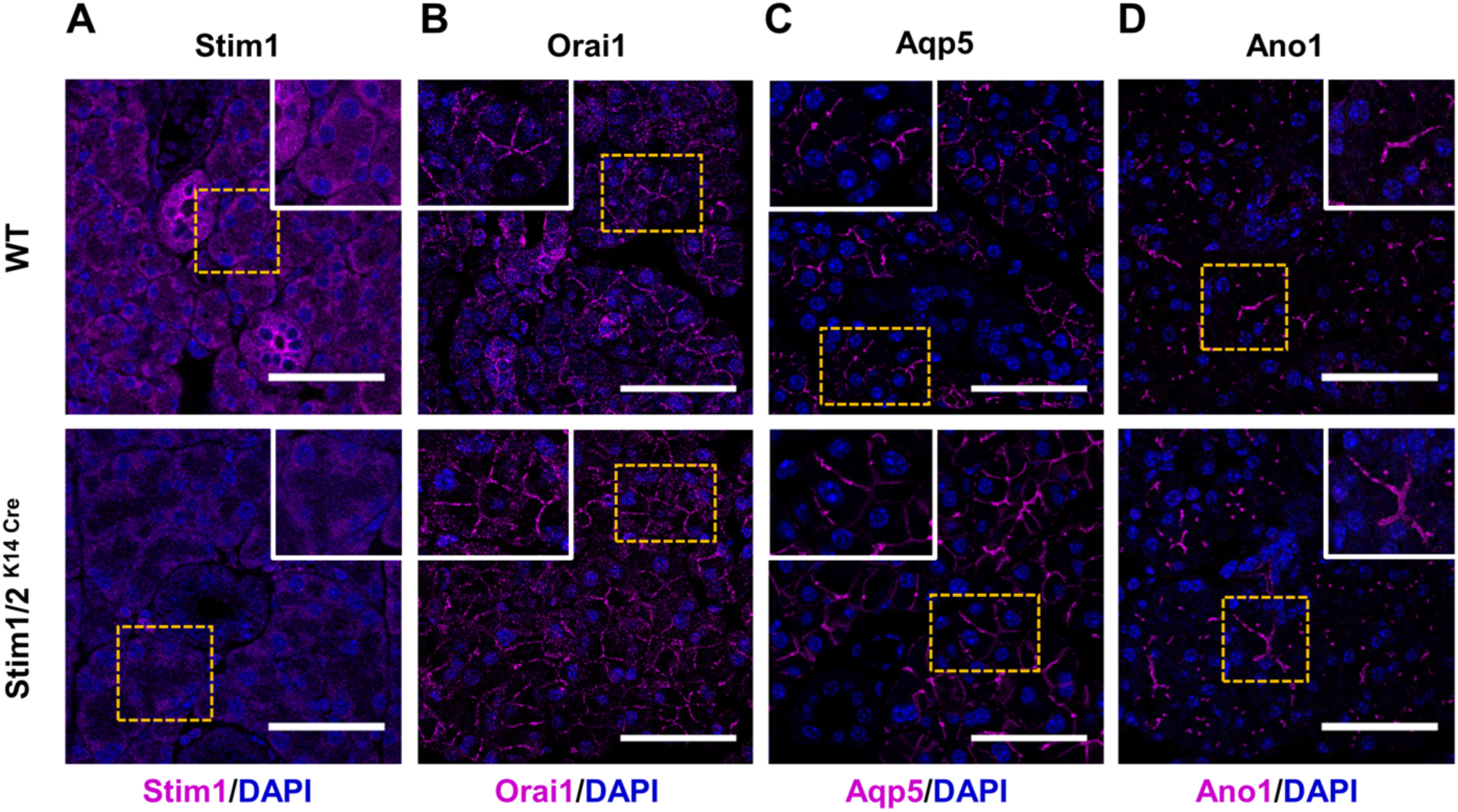
Subcellular localization of STIM1, ORAI1, AQP5 and ANO1 in parotid glands. **(A)** Representative images of Stim1 localization in parotid gland tissue sections of WT (top) and *Stim1/2^K14cre^* mice (bottom). **(B)** Localization of Orai1 in parotid gland tissue sections of WT (top) and *Stim1/2^K14cre^* mice (bottom). **(C)** Localization of Aqp5 in parotid gland tissue sections of WT (top) and *Stim1/2^K14cre^* mice (bottom). **(D)** Localization of Ano1 in parotid gland tissue sections of WT (top) and *Stim1/2^K14cre^* mice (bottom). In all images nuclear staining is highlighted by DAPI= blue. Inserts are higher magnification fields of areas highlighted by the squares. Scale bars = 50 µm.

### ANO1 function is reduced in parotid cells of *Stim1/2^K14Cre^* mice

CaCC are required for normal electrolyte and fluid secretion (35). Cl^-^ secretion into the lumen of the salivary glands generates a negative electrical potential drawing the paracellular transport of Na^+^ (16) and the subsequent transfer of water via the opening of AQP5 (36). The Cl^-^ channels ANO1 and BEST2 (bestrophin 2) are expressed in several secretory epithelia, including salivary glands, and are activated by intracellular Ca^2+^ (34, 37), but only ANO1 is essential for saliva production (25). Using whole cell patch clamp electrophysiology, we recently showed that ANO1 function was deficient in the human eccrine sweat gland cell line NCL-SG3 lacking *ORAI1* expression (27). Although ANO1 plays an essential role in salivary fluid secretion, it is not known if SOCE is required for the Ca^2+^ signal that activates ANO1 in salivary glands. To address this issue, we performed whole cell patch clamp electrophysiology in parotid cell preparations of WT and *Stim1/2^K14Cre^* mice as described (27). We found that ANO1 currents in parotid glands of WT mice evoked by depolarizing steps in the presence of 30 μM CCh are robust and sustained Cl^−^ currents (Figure 5A-5C). By contrast, parotid glands of *Stim1/2^K14Cre^* mice showed only transient CCh induced Cl^−^ currents which likely result from activation of ANO1 by Ca^2+^ released from ER Ca^2+^ pools (Figure 5A-5C). These data indicate that SOCE is required for the sustained activation of ANO1 in salivary glands.

**Figure 5.**
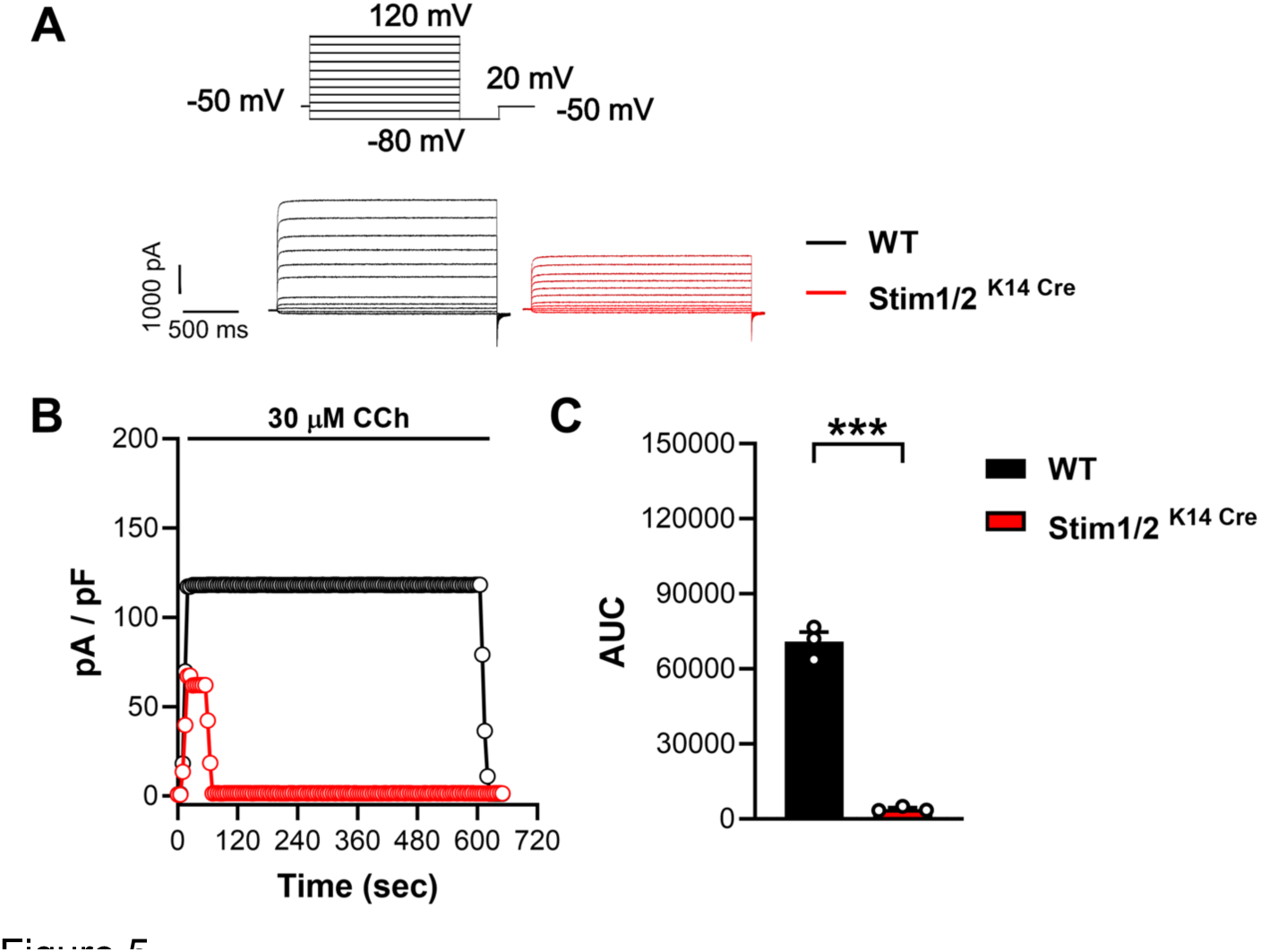
Sustained function of the Cl^-^ channel ANO1 is mediated by SOCE. Representative current traces of Cl^−^ currents measured by patch clamping in whole-cell configuration**. (A)** representative TMEM16 currents evoked by depolarizing steps shown in the insert in the presence of 30 uM CCh. **(B)** Time course of response to stimulation (at -80mV). **(C)** Area under the curve (AUC) from traces shown in (B). The AUC was integrated between 50 and 600 seconds (mean ± SEM of 3 independent experiments, n = 10 cells per experiment). Statistical significance was determined using a two-tailed Student’s t test. \*\*\**P* < 0.001.

### *Stim1/2^K14Cre^* mice do not show SjS-like phenotype

Because SOCE and ANO1 function are impaired in parotid glands of *Stim1/2^K14Cre^*mice, we next addressed whether these defects could be a driver for salivary gland inflammation and the onset of SjD. A diagnostic criterion of SjD is the finding of lymphocytic infiltrates, visually recognizable in salivary gland biopsies (5). We thus isolated parotid glands from 40-week-old female WT control and *Stim1/2^K14Cre^* mice and analyzed them using H&E staining. We found no clear alterations in the morphology of the salivary glands in *Stim1/2^K14Cre^* mice and no significant lymphocytic infiltration (Figure 6A and 6B). Immunofluorescence staining of parotid gland tissue sections with the T-lymphocyte marker CD3 showed no differences in CD3 expression between WT and *Stim1/2^K14Cre^* mice (Figure 6C). We next addressed whether the *Stim1/2^K14Cre^* mice showed elevated levels of autoantibodies commonly found in SjD patients. The classification criteria of SjD include the presence of anti-Ro (SSA) and anti-La (SSB) autoantibodies (5). While anti-Ro antibodies can also be associated with other autoimmune diseases such as systemic lupus erythematosus (SLE), rheumatoid arthritis and others, anti-La antibodies are more specific for SjD (7). We obtained serum from 40-week-old female *Stim1/2^K14Cre^* mice and WT controls and analyzed the presence of both anti-Ro and anti-La antibodies by ELISA. We found no significant changes in the levels of both autoantibodies in the serum of *Stim1/2^K14Cre^* mice (Figure 6D and 6E). These data suggest that *Stim1/2^K14Cre^* mice lack both cellular and humoral signs of autoimmune salivary gland inflammation indicative of SjD development.

**Figure 6.**
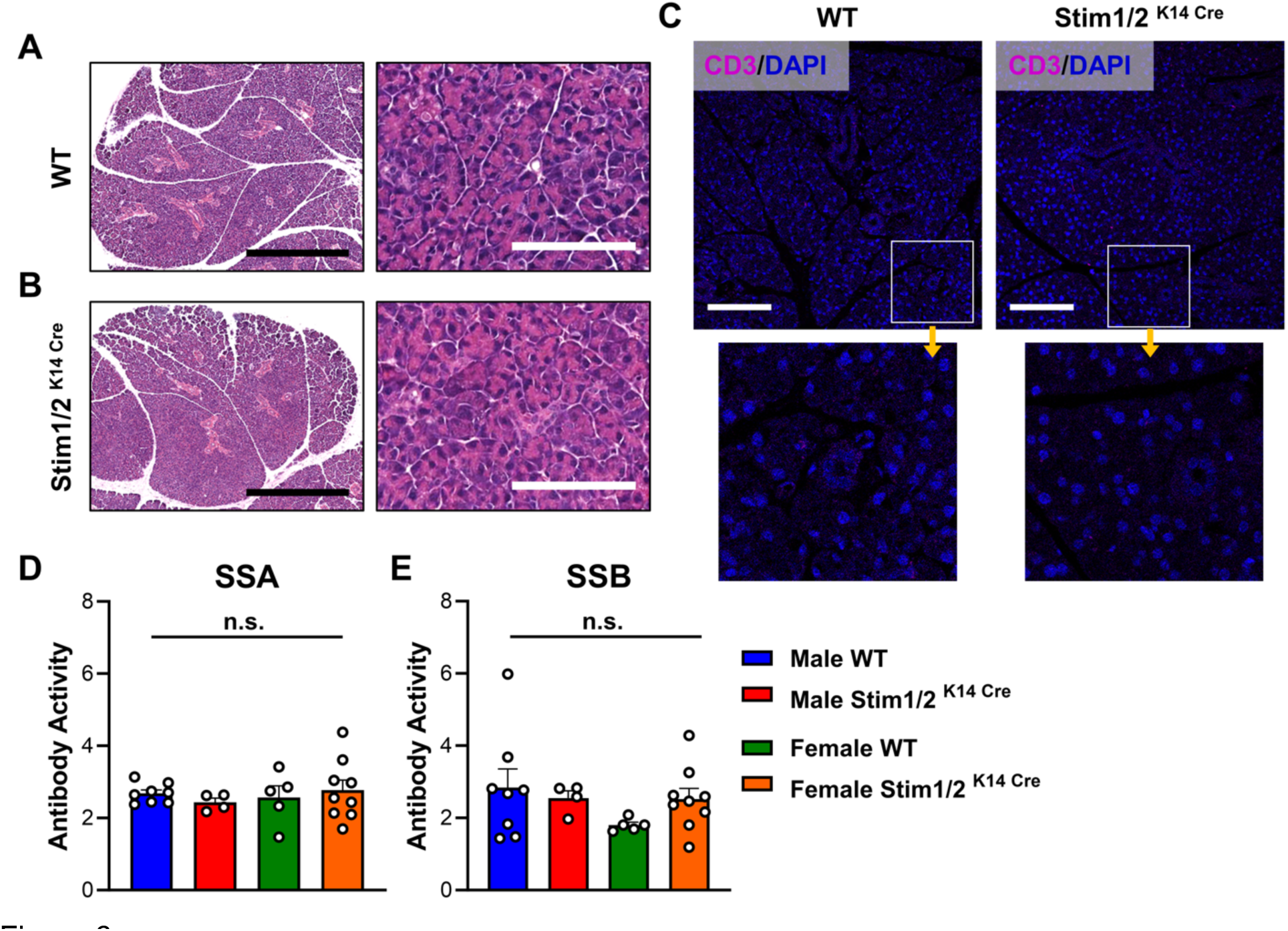
Loss of Stim1/2 in salivary glands does not induce disease. **(A and B)** Representative images of hematoxylin and eosin–stained sections of formalin-fixed, paraffin-embedded submandibular parotid glands of WT (n=3) and *Stim1/2^K14cre^* female mice (n=3). No mononuclear cells were identified. Scale: 500 µm (Left), 100 µm (Right) **(C)** Immunofluorescence staining of the T cell marker CD3 in WT and *Stim1/2^K14cre^* mice showing absence of CD3 in both. Inserts are higher magnification images of the regions highlighted by the squares. Scale: 100 µm **(D and E)** Quantification of the levels of SSA and SSB autoantibodies by ELISA in 40-week-old male and female WT and *Stim1/2^K14cre^* mice. N= at least 4 mice/group. One-way ANOVA followed by Bonferroni. n.s. = not significant.

### Activation of IFN signaling by DMXAA does not decrease saliva in *Stim1/2^K14Cre^* mice

Previous studies have shown that type I IFN responses are associated with the pathogenesis of SjD (38–41). In addition, the induction of IFN production in salivary glands by stimulation with dimethylxanthenone-4-acetic acid (DMXAA) elicits a SjD phenotype in mice including increased cytokine production and salivary gland hypofunction (39). DMXAA activates STING (stimulator of interferon genes) which is a mediator of type I IFN production (42). To address if hyposalivation renders *Stim1/2^K14Cre^*mice more susceptible to developing SjD-like disease in response to type I IFN production, we injected 17-week-old female *Stim1/2^K14Cre^* mice and WT littermates with DMXAA or vehicle to analyze salivary gland function (Figure 7A). The expression of the *Mx1* gene, which is induced by type I IFN (43), was upregulated in the salivary glands of both WT and *Stim1/2^K14Cre^* mice treated with DMXAA supporting the activation of type I IFN signaling (Figure 7B). We found that the volume of pilocarpine stimulated saliva production was reduced by approximately 28% in WT control female mice treated with DMXAA compared to WT mice injected with vehicle alone (Figure 7C) similar to the effects on saliva production reported previously (39). By contrast, female *Stim1/2^K14Cre^* mice treated with vehicle control had reduced saliva production which was not further reduced after DMXAA treatment (Figure 7C-7E). The lack of further hyposalivation in *Stim1/2^K14Cre^* mice treated with DMXAA might demonstrates that salivary gland dysfunction due to abolished SOCE does not render *Stim1/2^K14Cre^* mice more susceptible to exacerbation of SjD-like disease.

**Figure 7.**
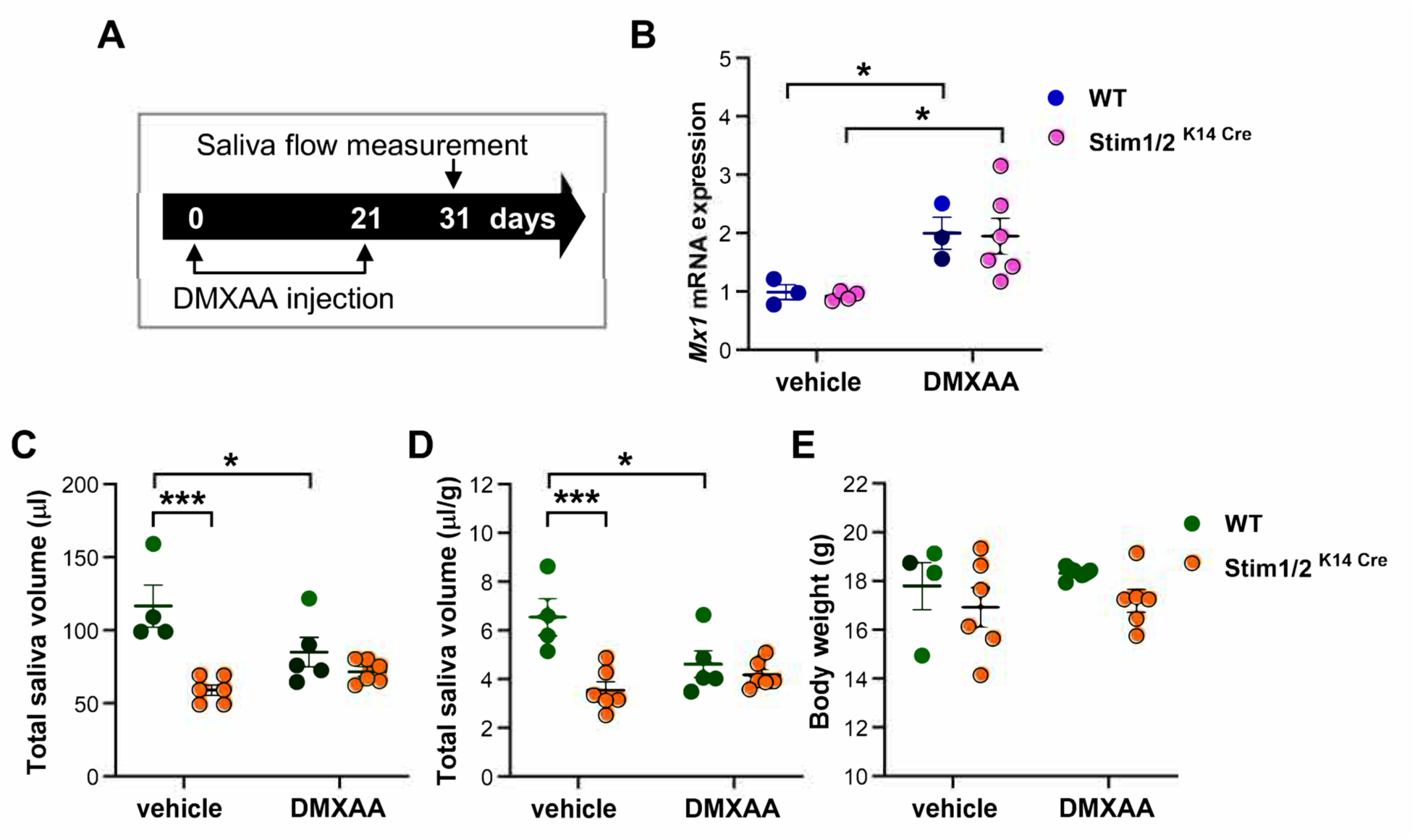
DMXAA treatment does not decrease saliva in *Stim1/2^K14cre^* mice. **(A)** Schematic of the DMXAA (dimethylxanthenone-4-acetic acid) treatment. **(B)** q-RT-PCR analysis of salivary glands of DMXAA treated and control mice showing upregulation in the expression of the proinflammatory cytokine *Mx1.* **(C)** Measurements of total volume of pilocarpine-induced saliva in vehicle- and DMXAA-injected groups of female mice. DMXAA-injected WT mice show a reduction in saliva flow but not *Stim1/2^K14cre^* mice. **(D)** Saliva volume corrected by body weight. **(E)** Body weight. Data represent the mean ±SEM from the analysis of n = 3 WT and n= 5 *Stim1/2^K14cre^* mice. (\**P* < 0.5, \*\*\**P* < 0.001, Two-way ANOVA followed by Tukey)

### Altered gene expression of immunity-associated genes

Genes in immunological pathways were among the top genes identified by IPA as being differentially expressed, which included complement system, phagosome formation and immune cell trafficking, all of which were downregulated in parotid glands of *Stim1/2^K14Cre^* mice (Figure 8A and Supplemental Figure 3). This included a downregulation of IFN-γ as an upstream regulator of dysregulated signaling pathways. This general downregulation of immunological pathways might be indicative of decreased inflammatory cascades and may have also affected the lack of further saliva secretion in the DMXAA treatment. Among the most downregulated immune-related genes in parotid glands of *Stim1/2^K14Cre^* mice identified by IPA were the toll-like receptor 8 (*Tlr8*) (p < 0.0001) and interferon regulatory factor 7 (*Irf7*) (Figure 8B). TLR8 is an intracellular single stranded RNA sensor found in endolysosomal membranes (44), which is involved in NF-kappaB signaling (45) upstream of IRF7 (46, 47), and has been implicated in SjD (48). Interestingly, it has been reported that knockdown of *STIM1* decreases TLR signaling in macrophages (49). The expression of *TLR8* mRNA was reported previously in whole preparations of salivary glands (50, 51), but it is unclear whether TLR8 is in fact expressed in acinar cells. To address this, we analyzed TLR8 localization by immunofluorescence in parotid glands of WT and *Stim1/2^K14Cre^* mice. We found that parotid acinar cells of WT mice had strong intracellular TLR8 staining compared to *Stim1/2^K14Cre^*mice (Figure 8C and 8D) which showed lower fluorescence intensity (Figure 8E). TLR8 was absent in ductal cells (Figure 8C and 8D). This decrease in *Tlr8* mRNA and protein expression in *Stim1/2^K14Cre^* mice suggested a possible link between SOCE and TLR8. To further address this, we analyzed the expression of *TLR8* and *TLR7*, which have been implicated in SjD (48), in two human salivary gland cell lines; a ductal and an acinar cell model shared by Dr. Jay Chiorini (NIDCR). The acinar cells showed significantly higher expression of both genes than the ductal cells (Supplemental Figure 4), similar to the results we obtained in mouse salivary glands where TLR8 was only visible in salivary gland acini. We then stimulated SOCE in the human acinar salivary gland cell line with CPA (20 µM) and analyzed the expression of *TLR8*, *TLR7* and *IRF7* by qRT-PCR. CPA treatment did not induce endoplasmic reticulum stress (Supplemental Figure 5). We found that SOCE activation significantly upregulates the expression of all three genes (Figures 8E,8G) supporting a link between SOCE and TLR expression.

**Figure 8.**
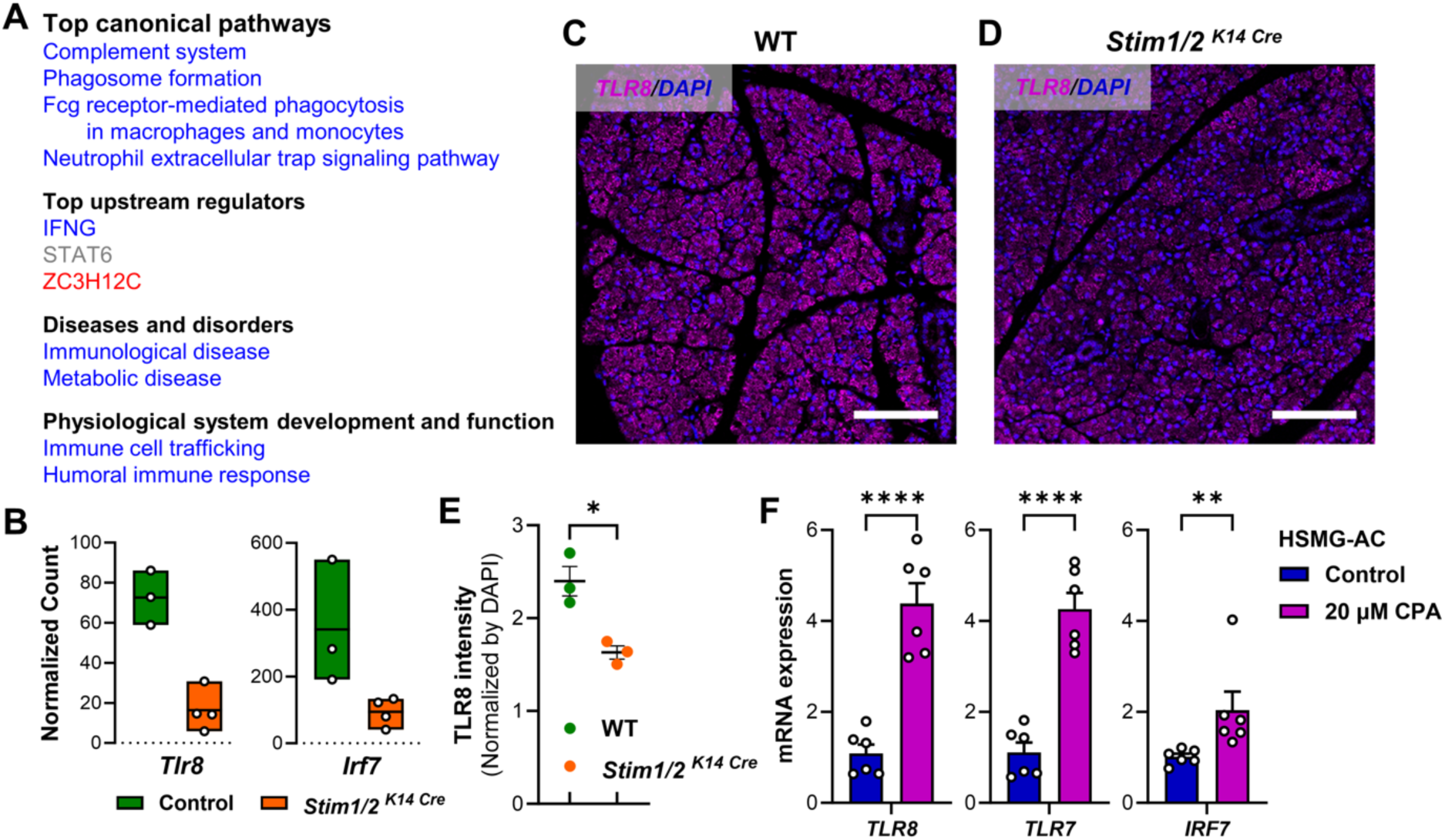
RNASeq shows decreased inflammatory pathways in *Stim1/2^K14cre^* mice. **(A)** Top canonical pathways and others highlighted by IPA in the RNASeq of parotid glands in WT and *Stim1/2^K14cre^* mice. Blue = downregulated, red = upregulated. **(B)** Normalized counts of *Tlr8* and its downstream target *Irf7* analyzed in the RNASeq. n = 3 WT, n = 4 *Stim1/2^K14cre^* mice. **(C and D)** Representative immunofluorescence staining of TLR8 in acinar cells of parotid gland tissue sections of WT and *Stim1/2^K14cre^*mice. Nuclear staining DAPI (blue). **(E)** Densitometric quantification of TLR8 fluorescence in tissue sections of WT and *Stim1/2^K14cre^* mice. n = 3 sections/mice per group. **(F)** q-RT-PCR analysis of *TLR8*, *TLR7*, and *IRF7* genes in the human submandibular acinar cells (HSMG-AC) untreated and in cells stimulated with 20 µM CPA for 2 hours. Data represent the mean ±SEM from at least 3 different experiments (\**P* < 0.5, \*\**P* < 0.01, \*\*\*\**P* < 0.0001. Un-paired Student’s *t*-test).

### STIM1 and TLR8 gene expression are elevated in SjS biopsies

It is currently unknown if the molecular components of SOCE are altered in SjD. We obtained minor salivary glands (MSG) biopsies and tissues sections from SjD patients and symptomatic non-SjD controls from the Sjogren’s International Collaborative Clinical Alliance (SICCA) consortium. Determination of positive SjD status was made based on the ACR-EULAR Classification Criteria for SjD scored from 1-9 (52) (Supplemental Table 1). Samples were divided into symptomatic non-SjD controls with scores of ≤ 2, SjD patients with intermediate scores of 4-5, and patients with higher scores of 6-9 (Figure 9A). We analyzed the gene expression levels of *STIM1*, *ORAI1*, *IR3R2, IP3R3* and *AQP5* by qRT-PCR. We found that *STIM1* expression was elevated in MSG samples of SjD patients with high scores of 6-9 but not in SjD patients with lower scores of 4-5 compared to controls (Figure 9B). *ORAI1* expression did not change in SjD patients compared to controls (Figure 9B). Both *IP3R2* and *IP3R3* levels were upregulated in SjD samples with higher scores (Figure 9B). Data on the expression of *AQP5* in SjD patients has been conflicting in the past. We found that the expression of *AQP5* was significantly decreased only in SjD patients with high scores compared to controls (Figure 9B). Moreover, we analyzed the localization of STIM1, ORAI1 and AQP5 proteins in tissue sections of controls and SjD patients. Although the SjD samples exhibited changes in glandular structure because of lymphocytic infiltrates (Supplemental Figure 6A), there were no obvious changes in the localization of STIM1 in SjD compared to control samples (Figure 9C and 9D). Of note, STIM1 showed stronger signals in salivary gland ductal cells compared to acinar cells in both control and SjD samples (Figure 9C and 9D). AQP5 was expressed at the apical surface of acinar cells without appreciable differences between control and SjD samples (Figure 9C and 9D). The expression and localization of ORAI1 were similar in control and SjD tissues showing a strong lateral plasma and some basal membrane signals but showed no overlap with the apical localization of AQP5 using two separate antibodies (Figure 9E and 9F, and Supplemental Figure 6B). These data suggest that the cellular localization of STIM1, ORAI1 and AQP5 in salivary gland cells does not change in SjD patients compared to non-SjD controls. Because *STIM1* mRNA levels were increased in MSG samples of SjD patients (Figure 9B) and *Stim1/2^K14Cre^* mice had lower levels of *Tlr8* in their parotid glands (Figure 8B and 8E), we analyzed the expression of *TLR8* and *TLR7* by qRT-PCR in MSG biopsies of SjD patients. We found that mRNA levels of both genes were significantly elevated with an average fold increase of ≥ 100 in SjD patients (Figure 9G) suggesting that STIM1 regulates TLR expression in SjD.

**Figure 9.**
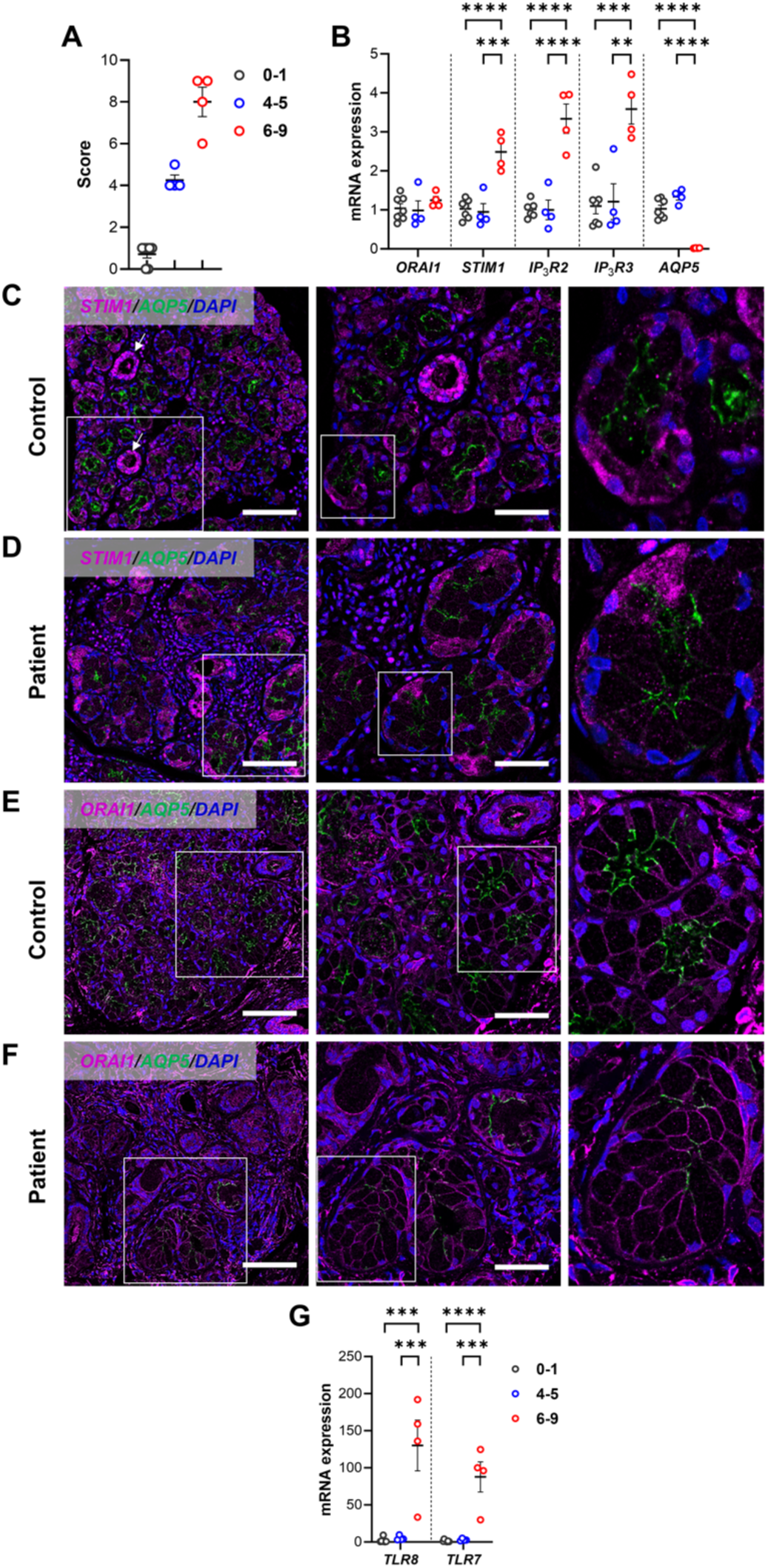
Gene expression changes in SjD patients. **(A)** Distribution of the patient’s scores obtained from SICCA using the 2016 ACR-EULAR system. Controls: score 0-1 (n = 7), SjD patient: score 4-5 (n = 4), SjD patients: score 6-9 (n = 4). **(B)** q-RT-PCR analysis of *ORAI1*, *STIM1*, *IP_3_R2*, *IP_3_R3* and *AQP5* genes in minor salivary gland (MSG) biopsies from the SICCA samples. **(C and D)** Subcellular localization of STIM1 and AQP5 in fixed tissue sections of minor salivary glands of healthy (C) and SjD patients (D). Arrows indicate ductal cells. White squares indicate regions enlarged from left to right. **(E and F)** Subcellular localization of ORAI1 and AQP5 in fixed tissue sections of minor salivary glands of healthy (E) and SjD patients (F). White squares indicate regions enlarged from left to right. N=3 samples per group were analyzed. Scales: 100 µm (Left), 50 µm (Right) **(G)** q-RT-PCR analysis of *TLR8* and *TLR7* genes in MSG biopsies from the SICCA samples. Data (B) and (G) represent the mean ±SEM from each sample run in triplicates. (\*\**P* < 0.01, \*\*\**P* < 0.001, \*\*\*\**P* < 0.0001, Two-way ANOVA followed by Tukey).

## Discussion

Ca^2+^ signaling is important for saliva production and has been implicated in SjD (13, 53). However, the broader mechanisms mediating the loss of salivary gland function in SjD and whether Ca^2+^ signaling is involved in SjD development remains unclear (7). Salivary fluid production is a complex process initiated by parasympathetic input into the glands (15) activating muscarinic acetylcholine receptors (18). Two main sources of cytosolic Ca^2+^ contribute to the signaling cascade in acinar cells, the more transient IP_3_R-mediated release of Ca^2+^ from ER stores (32), and the sustained Ca^2+^ influx via SOCE(11).

The role of IP_3_R function has been studied in vivo and two isoforms are considered essential in salivary glands, type 2 (IP_3_R2) and type 3 (IP_3_R3) (32). Double homozygous mouse mutants of *Ip3r2* and *Ip3r3* showed abnormal Ca^2+^ signaling and a deficit in saliva secretion (32). Further, salivary glands of SjD patients showed decreased expression of both IP_3_R subtypes and acinar cells displayed decreased ER Ca^2+^ release consistent with abnormal Ca^2+^ signaling in SjD patients, suggesting that a deficit in IP_3_R signaling underlies hypofunction of the glands in SjD (53). The role of SOCE in salivary glands has been addressed by siRNA knockdown of *STIM1* and *ORAI1* genes and the ORAI1 mutant pore R91W in human salivary gland cell lines which showed decreased Ca^2+^ transients (54, 55). However, how the loss of the molecular components of SOCE affects salivary gland function (11) in vivo is poorly understood, and there are no reports of salivary gland function in mice lacking *Stim1* or *Orai1*. By contrast, the role of the TRPC1 channel in salivary glands is better known. *Trpc1^-/-^* mice were reported to have reduced SOCE and decreased salivary gland fluid secretion (22), and the current interpretation suggest that SOCE-mediated Ca^2+^ entry triggers the recruitment of TPRC to the plasma membrane, being a separate channel from ORAI1, but influencing SOCE (11, 54).

Because the function of SOCE in salivary glands has been largely studied using salivary gland cell lines, here we analyzed the effects of *Stim1* and *Stim2* deletion in salivary glands in vivo. We found that stimulated saliva production in *Stim1/2^K14Cre^* mice using pilocarpine resulted in a significant decrease in saliva flow in both male and female mice. Moreover, parotid cell preparations loaded with Fura 2-AM and stimulated with CPA to activate SOCE showed robust Ca^2+^ transients in acinar cells of WT male and female mice, but SOCE was nearly abolished in the parotid glands of *Stim1/2^K14Cre^* mice of both sexes, suggesting that loss of *Stim1/2* in salivary glands affected saliva fluid production because of ablated SOCE. Physiologically, a cytosolic Ca^2+^ increase is required for the activation of Ca^2+^ activated Cl^-^ channels that control normal electrolyte and fluid secretion (35, 56). Cl^-^ is a key element in the transepithelial transport of electrolytes and fluid secretion. The electroneutral Na^+^/K^+^/2Cl^-^ transporter at the basolateral membrane of acinar cells plays an important role in generating an increase in intracellular Cl^-^ concentrations five times above its electrochemical gradient (16). Cl^-^ efflux and its luminal accumulation generates a transepithelial osmotic gradient driving the movement of water via AQP channels (16). cAMP activated CFTR (cystic fibrosis transmembrane conductance regulator) conducts Cl^-^ in some epithelial cells such as pancreatic ducts, airway cells and others, whereas Ca^2+^ activated Cl^-^ channels secrete Cl^-^ in acinar cells of the pancreas, salivary glands, and sweat glands (57). In salivary glands, the main CaCC required for salivary fluid secretion is ANO1 (TMEM16A) (25, 34). Recent cryo-EM studies revealed that ANO1 is activated by the binding of Ca^2+^ to a site in the vicinity of the channel pore triggering a conformational rearrangement and the opening of the pore (58). However, whether SOCE activates ANO1 is salivary glands is unknown. To address this, we performed whole-cell patch clamp electrophysiology and showed that parotid acinar cell of *Stim1/2^K14Cre^* mice lacked sustained CCh-induced Cl^-^ currents, indicating a critical role of SOCE in ANO1 activation in salivary glands, similar to the role of SOCE in eccrine sweat glands (27). Although there are no reports on salivary fluid secretion in *Orai1^-/-^* mice, a previous study showed loss of lacrimal gland function in *Orai1^-/-^* mice (59), and we showed decreased sweat production in *Orai1^K14Cre^* (27). This data highlights an important role on SOCE in mediating exocrine gland function and epithelial Cl^-^ transport.

Our bulk RNASeq analysis of parotid glands *Stim1/2^K14Cre^*did not show differential expression of genes involved Ca^2+^ signaling pathways that mediate saliva flow, with the exception of *Stim1* and *Stim2*, which were significantly downregulated. Our qRT-PCR and Western blot data further demonstrate that expression levels of *Orai1*, *Ip3r1*, *Ip3r2*, *M3r*, *Aqp5* and *Ano1* are comparable in *Stim1/2^K14Cre^*and WT mice (Figure 3 and Supplemental Figure 2). Previous studies have reported that ORAI1 is localized at the apical and lateral poles in acinar cells of submandibular glands (54), similar to its localization in acinar cells of another exocrine gland, the pancreas (29). We here show that ORAI1 is located in the lateral membrane in the human MSG. STIM1 fluorescence was strong in ductal cells in WT mice and was absent in tissues of *Stim1/2^K14Cre^* mice. Similarly, ANO1 localization at the apical pole appeared unchanged in salivary glands of *Stim1/2^K14Cre^* mice. AQP5 is an important channel involved in water transport in salivary glands that is found at the apical pole of salivary acini (33). The role of AQP5 in salivary glands is supported by a 60% reduction of pilocarpine stimulated saliva in *Aqp5* transgenic mice (60). In our study, we found no differences in the localization of AQP5 between WT and *Stim1/2^K14Cre^* mice although Ca^2+^ signals were reported to be important for the recruitment of AQP5 to the plasma membrane (61).

Clinical hallmarks SjD include the infiltration of T and B lymphocytes in salivary glands causing inflammation, the presence of anti-SSA/Ro and anti-SSB/La antibodies, and hypofunction of salivary and lacrimal glands (5, 7, 9, 62). Although ageing ∼40 weeks old *Stim1/2^K14Cre^* mice *showed* hyposalivation and altered Ca^2+^ signaling in their salivary glands, the histological analysis of parotid glands did not show evidence of lymphocytic infiltrates. Similarly, the analysis of parotid glands by immunofluorescence failed to show increased infiltration by CD3^+^ T cells in the glands of Stim*1/2^K14Cre^* mice. Further, the serum levels of anti-SSA/Ro and anti-SSB/La antibodies were not elevated in *Stim1/2^K14Cre^* mice compared to littermate controls. Combined, these data demonstrate that impaired saliva production and Ca^2+^ signaling in the parotid glands of *Stim1/2^K14Cre^*mice are not associated with salivary gland inflammation or the production of autoantibodies. Moreover, the induction of a STING-dependent type I IFN response by injection of mice with DMXAA did not further exacerbate the salivary gland dysfunction of *Stim1/2^K14Cre^* mice. On the contrary, our RNASeq analysis demonstrated an overall decrease in inflammatory pathways including IFN and *Tlr8* signaling. Collectively, these data indicate that loss of SOCE in salivary gland acinar cells suppresses inflammatory responses rather than rendering *Stim1/2^K14Cre^* mice susceptible to salivary gland inflammation and autoantibody production.

TLRs play a central role in the initiation of innate immune responses to pathogens (44, 63, 64). The expression of several members of the TLR family have been previously reported in salivary glands (50, 51, 65). Of these, TLR7 and TLR8 have been implicated in SjD (48, 66). *Tlr8* KO mice showed SjD-like phenotypes including high levels of SSA and SSB autoantibodies, sialadenitis, ectopic lymphoid structures and elevated cytokine levels (48). These effects of *Tlr8* deletion were abrogated in mice lacking both *Tlr7* and *Tlr8* expression, suggesting that the effects of TLR8 on suppressing SjD-like disease in mice are dependent on TLR7 (48). Interestingly, transgenic mice harboring high copy numbers of human TLR8 showed an inflammatory phenotype that was restricted to exocrine tissues including salivary glands (67). A gain-of-function mutation in *TLR7* was recently shown to cause SLE (68), which, like SjD, is an autoimmune disease associated with an increased type I IFN response and can result in secondary Sjoegren’s syndrome. Our RNASeq analysis of salivary glands from WT and *Stim1/2^K14Cre^* mice showed that whereas *Tlr7* was not differentially expressed, *Tlr8* was one of the most downregulated genes in *Stim1/2^K14Cre^* mice, suggesting a connection between SOCE and TLR8 expression. We provide further evidence for a connection between SOCE and the expression of TLRs because induction of SOCE in a human salivary gland cell line resulted in a significant upregulation of *TLR8, TLR7* and *IRF7* gene expression. *IRF7* specifically is considered as an important regulator of IFN-dependent immune responses which may have influenced the lack of IFN in DMXAA treated mice (69). These findings are consistent with a previous study in macrophages, which showed that TLR mediated cytokine signaling was potentiated by extracellular Ca^2+^ whereas knockdown of *STIM1* decreased the TLR response (49). TLR8 and TLR7 are localized in endolysosomes and their activation depends on the maintenance of an acidic compartment because TLR8 responses are suppressed when lysosomal acidification is prevented by bafilomycin A (44, 70). Interestingly, there is evidence that lysosomal Ca^2+^ is modulated by STIM1 (71). Taken together, our data suggest that SOCE regulates the expression of TLR8 and that abolishing SOCE in *Stim1/2^K14Cre^*mice may suppress TLR8 mediated salivary gland inflammation.

To address possible changes in the expression of molecular components of the SOCE pathway in human patients diagnosed with SjD, we measured the levels of *STIM1* and *ORAI1* mRNA and protein in salivary gland biopsies. Interestingly, the expression levels of *STIM1* were significantly increased in SjD patients with high disease scores, whereas the localization of STIM1 protein determined by immunofluorescence was similar in patients and controls. By contrast, *AQP5* mRNA levels were significantly decreased in patients with high disease scores, consistent with a previous report from a SjD patient cohort from Norway analyzed by immunohistochemistry (13). Another report found that AQP5 levels were unchanged in SjD patients (53), which might be explained by the inclusion of patients with relatively low disease scores. Our analysis showed strong differences in *AQP5* gene expression only in SjD patients with high disease scores but not in those with intermediate scores. It is noteworthy that *ORAI1* expression levels were unchanged in salivary glands of SjD patients and that the localization of ORAI1, STIM1 and AQP5 was similar in SjD patients and non-SjD controls was similar.

Collectively, our data show that loss of *Stim1* and *Stim2* in mouse salivary glands results in hyposalivation caused by the loss of SOCE and impaired Ca^2+^ dependent activation of the Cl^-^ channel ANO1. This defect in Ca^2+^ signaling did not, however, induce salivary gland inflammation or the production of autoantibodies associated with SjD in human patients. We hypothesize that the absence of salivary gland inflammation is due to a decrease in TLR8 expression and signaling.

## Acknowledgements

This work was funded by National Institute of Dental and Craniofacial Research (NIDCR) grants DE027981 to R.S.L and S.F., and DE014756 to D.I.Y. Human samples used in this study are from the Sjögren’s International Collaborative Clinical Alliance (SICCA) Next Generation Studies, funded under contract # 1U01DE028891-01A1 (and previously under N01 DE-32636 and #HHSN26S201300057C) by the National Institute of Dental and Craniofacial Research. The content and findings of this manuscript does not necessarily reflect the opinions or views of the SICCA investigators, the NIH or NIDCR. We thank Dr. Jay Chiorini (NIDCR) for sharing the salivary gland cell lines. We also thank Dr. Veronica Costiniti for assistance with immunofluorescence experiments. We than Dr. Lily Jan at Howard Hughes Medical Institute, UCSF, for the ANO1 antibody.

## Materials and methods

### Human samples

Deidentified human minor salivary gland biopsies and tissue sections were obtained from the Sjogren’s International Collaborative Clinical Alliance (SICCA) consortium. The classification of the tissue samples with focus score is shown in Table S1.

### Mice

Wild-type and *Stim1/2^K14^ ^Cre^* mice were reported previously (27). All mice were maintained on a C57BL/6 genetic background and used between 6 and 40 weeks of age. Animals were kept in specific-pathogen-free rooms in the vivarium of NYU College of Dentistry in normal light-dark cycles under the constant temperature of 22 ± 2°C and 60 ± 10% humidity. Animal studies were approved under the Institutional Animal Care and Use Committee (IACUC) protocol # IA16-00625.

### Saliva secretion measurements

Saliva secretion was measured in mice every month from 9-to 39-weeks of age. Body weight was measured prior to anesthetize the mice. Mice were anesthetized with a cocktail of ketamine (0.06 mg/g) and xylazine (0.008 mg/g) diluted in 0.9% saline by intramuscular injection. Saliva was collected for 20 min following pilocarpine (0.5 mg/kg) (P6503, Sigma) stimulation subcutaneously as described (72).

### Primary culture of parotid gland cells

Parotid glands were excised, and connective and fat tissues were removed. Parotid glands were finely minced and digested in Dulbecco’s modified eagle medium (DMEM, 11960-044, Gibco) containing 0.25 mg/ml of Liberase (TL, 05401020001, Roche) and 1 % BSA (A7888, Sigma) for 25 min. The digested tissues were spun down (2000 rpm, 5 min) and cell pellets were washed and re-suspended in DMEM with 1 % BSA. The digestion medium and parotid gland cells were continuously gassed with a mixture of 95 % O_2_ and 5 % CO_2_ and incubated in a shaker/water bath at 37 °C until use.

### Cell Culture

Human submandibular acinar cells (NS-SV-AC) and ductal cells (NS-SV-DC) were obtained from Dr. John Chiorini at National Institute of Dental and Craniofacial Research (NIDCR). Acinar cells were cultured in Defined keratinocyte-SFM (10744019, Gibco), while ductal cells were cultured in KGM™ Gold Keratinocyte Growth Medium BulletKit (00192060, Lonza). Cells were harvested by treating 0.05 % Trypsin/EDTA (25300062, Gibco), and washed with Dulbecco’s modified eagle medium (DMEM, 12430054, Gibco) supplemented with 10% fetal bovine serum (FBS, Gibco, #26140-079). Cells were subcultured at a 1:5 ratio (for acinar cells) or a 1:2 ratio (for ductal cells) in 100 mm dishes, maintaining them at 37°C in a humidified 20 % O_2_/ 5 % CO_2_ incubator for 4-5 days.

### Cytosolic Ca^2+^ measurements

Primary cultured parotid gland cells were loaded with Fura 2-AM (F1221, Invitrogen) (10 μg/ml) in DMEM with 1 % BSA and incubated in the shaker/water bath at 37 °C for 45 min under dark conditions. Cells were continuously gassed with a mixture of 95 % O_2_ and 5 % CO_2_ during incubation. Cells were washed 3 times with normal Ringer’s solution (pH 7.4) as follows (mM): 150 NaCl, 4.5 KCl, 2 CaCl_2_, 1 MgCl_2_, 10 D-glucose, and 5 HEPES (Sigma-Aldrich, USA), and kept on ice. Cells were allowed to attach to Cell-Tak (354240, Corning)-coated glass cover slips for 10 min. Cytosolic Ca^2+^ concentration ([Ca^2+^]_cyto_) was measured first in Ca^2+^-free Ringer solution (pH 7.4) [150 NaCl, 4.5 KCl, 0 CaCl_2_, 3 MgCl_2_, 10 D-glucose, and 5 HEPES (mM)] for 1 min followed by stimulation with 30 µM cyclopiazonic acid (CPA, Sigma, #1530) in Ca^2+^-free Ringer solution to deplete ER store, then re-addition of 5 mM Ca^2+^-containing normal Ringer solution to stimulate Ca^2+^ influx by store-operated Ca^2+^ entry (SOCE). During the [Ca^2+^]_cyto_ measurements, cells were continuously perfused with Ringer’s solution by a perfusion system (VC-6/8 valve controller) at 4 ml/min which was controlled by electrical control valves (Harvard Bioscience Inc., USA). Fura 2-AM fluorescence was measured using a Nikon Ti2-E Eclipse microscope (Nikon) with an objective Nikon S Fluor x20 and a DS-Qi2 digital SLR camera (Nikon) controlled by NIS elements software (Version 5.20.01, Nikon). Cells were excited at 340 and 380 nm with an emission of 510 nm. Images were acquired every 3 seconds, and 340/380 ratio was calculated.

### CPA treatment

Human submandibular acinar cells were treated with various µM concentrations of CPA (1, 5, 10, 15, 20, 25) for 2 hrs to first determine if CPA induced ER stress by analyzing *GRP78* mRNA levels. 20 μM did not induce ER stress and was then used to stimulate SOCE in the acinar cell lines. Total RNA was isolated from untreated and stimulated cells.

### RNA isolation and real-time PCR

Total RNA was isolated using miRNeasy Mini Kit (1038703, Qiagen) for RNASeq and RNeasy mini kit (74106, Qiagen). A NanoDrop spectrophotometer (13400519, Thermo Scientific) was used to measure the 260/280 ratio. RT was performed using the iScript cDNA Synthesis Kit (1708891, Bio-Rad) in a thermal cycler (T100, Bio-Rad) following manufacturer’s instructions. For qPCR, we used the Powerup SYBR Green master mix (A25742, Applied Biosystems) according to manufacturer’s protocol and performed the experiments in a thermal cycler (CFX, Bio-Rad). Gene expression levels were evaluated by the 2^-(ΔCT)^ or 2^-(ΔΔCT)^ value, which was normalized by the housekeeping gene, GAPDH. All primers were used at the final concentration 0.25 µM. All primer sequences used in this study can be found in Supplemental Table 2.

### Patch clamp electrophysiology

For measurements of Cl^−^ currents, parotid acinar cells were acutely isolated and allowed to adhere to Cell-tak coated glass coverslips for 30 minutes prior to experimentation and then transferred to a chamber containing extracellular bath solution (155 mM tetraethylammonium chloride to block K^+^ channels, 2 mM CaCl2, 1 mM MgCl2, 10 mM HEPES, pH 7.2). Cl^−^ currents in individual cells were measured in the whole cell patch clamp configuration using pClamp 9 and an Axopatch 200B amplifier (Molecular Devices). Recordings were sampled at 2 kHz and filtered at 1 kHz. Pipette resistances were 3–5 megaohms, and seal resistances were greater than 1 gigaohm. Pipette solutions (pH 7.2) contained 60 mM tetraethylammonium chloride, 90 mM tetraethylammonium glutamate, 10 mM HEPES, and either 1 mM EGTA without added Ca^2+^ to yield negligible free Ca^2+^ or 5 mM EGTA with 4.3 mM CaCl2 added to yield 1 μM free Ca^2+^. Alternatively, 1 mM HEDTA (N-(2-hydroxyethyl)ethylenediamine-N,N’,N’-triacetic acid) and 20 μM CaCl2 were used in the internal pipette solution to better mimic physiological buffering and basal [Ca^2+^]_cyt_ conditions (∼100 nM Ca^2+^). Free [Ca^2+^] was estimated using Maxchelator freeware (http://maxchelator.stanford.edu/). Agonists were directly perfused onto individual cells using a multibarrel perfusion pipette.

### Histology

Mice were sacrificed at 40-weeks old at the end of measuring saliva flow. Excised parotid glands were fixed in 4 % of paraformaldehyde (PFA) in phosphate-buffered saline (PBS) (J19943-K2, Thermo scientific) for 48 hours at 4 °C. After rinsing the fixed tissues with PBS, the tissues were embedded in paraffin and sectioned ∼5 μm thickness. Sectioned tissues were stained with hematoxylin and eosin (H&E). The images of the H&E stained pathology slides were taken using a digital pathology slide scanner (Leica, Aperio CS2) and analyzed by one of us (Dr. Fang Zhou, a clinical pathologist).

### Immunofluorescence (IF)

For IF, unstained paraffin tissue sections were de-paraffinized using xylene (UN1307, Fisher scientific) followed by rehydration with different % of ethanol (From 100 % to 70 %, 111000190, Pharmco) and the antigen retrieval step with citric buffer (pH 6, C9999, Sigma) in heat for 20 min. Sections were then blocked with 5 % of skim milk (170-6404, Bio-Rad) in PBS-T (Containing 0.1 % of Tween-20) for 45 min at room temperature (22-25 °C) before applying primary antibodies overnight (4 °C). The following primary antibodies were used: Rabbit anti-Orai1 (1:50, from Dr. Stefan Feske) and mouse anti-ORAI1 (1:50, SAB3500126, SIGMA), rabbit anti-STIM1 (1:200, HPA012123, Sigma), rabbit anti-AQP5 (1:250, Ab92320, Abcam), and rabbit anti-ANO1 (or TMEM16A) (1:25, from Dr. Lily Jan at Howard Hughes Medical Institute at University of California San Francisco) in 5 % of skim milk in PBS-T. After washing in PBS-T, secondary antibodies were applied including Alexa flour 647 donkey anti-rabbit IgG (H+L) (1:200, A31573, Life technologies) and in 5 % of skim milk in PBS-T for 1 hour at room temperature. Sections were washed in and mounted using Prolong gold antifade reagent with DAPI (P36931, Invitrogen). Fluorescence was measured with using a Leica SP8 confocal microscope (Leica Microsystems) controlled by LAS X imaging software (Leica Microsystems). The acquired images were analyzed using imageJ software (National Institutes of Health).

### Western Blot Analysis

Parotid glands were excised, and the connective and fat tissues were removed. Tissue lysates were prepared using Pierce RIPA buffer (89900, Thermo Scientific) supplemented with Halt^TM^ phosphatase inhibitor cocktail (1861277, Thermo Scientific) though homogenization using a pestle and sonification prior to centrifugation at 10,000 rcf for 20 min at 4 °C. Proteins were quantitated using the Pierce^TM^ BCA Protein Assay (23227, Thermo Scientific). The protein samples were denatured using 1x Laemmli sample buffer with 2.5% 2-mercaptoethanol (M3148, Sigma) for each sample at 95 °C for 5 min (STIM1) and 70 °C for 10 min (ANO1 and AQP5). After denaturation, the samples were loaded at 40 µg/well into 10 % Mini-Protean TGX Gels (4561033, Bio-Rad) and separated by SDS-polyacrylamide gel electrophoresis before transferring onto Immun-Blot PVDF Membrane (162-0177, Bio-Rad). The membrane was block with Blotting-Grade Blocker (1706404, Bio-Rad) for 1 hr and then incubated with primary antibodies overnight at 4 °C. The primary antibodies included rabbit anti-STIM1 (1:1000, HPA012123, Sigma), rabbit anti-ANO1 (1:100, from Dr. Lily Jan), rabbit anti-AQP5 (1:1000, ab78486, Abcam) and rabbit anti-GAPDH (1:1000, 21185, Cell Signaling). Tris-buffered saline (1706435, Bio-Rad) with Tween 20 (1706531, Bio-Rad,) was used to wash the membranes before incubating with goat anti-rabbit horseradish peroxidase conjugated secondary antibody at 1:5000 dilution (1706515, Bio-Rad) for 90 min at room temperature. The signal was detected using Super Signal^TM^ West Pico PlUS Chemiluminescent Substrate (34577, Thermo Scientific).

### ELISA

Blood samples were collected 40-weeks old. Serum was separated by centrifugation at 2000 rpm for 10-15 min after incubation without anti-coagulant at 4 °C overnight, and then stored at -80 °C. To measure Sjögren’s Syndrome A (SSA) and Sjögren’s Syndrome B (SSB), we used the mouse anti-SSA (5710, Alph diagnostic international) and anti-SSB ELISA kits (5810, Alph diagnostic international) according to manufacturer’s instructions. The optical density of the standards and samples was recorded at 450 nm wavelength using a microplate reader (Promega, GloMax 9301). All samples were run in triplicates.

### DMXAA treatment

Female mice (12-to 16-weeks old) were injected subcutaneously with (20 mg/kg body weight) DMXAA (5601, Tocris) dissolved in 5% sodium bicarbonate solution. For long-term experiments, the mice received 2 injections (On the day 1 and day 21). The dose of DMXAA was based on previously published literature (39). Control mice were injected similarly with the vehicle alone. On the day 31, saliva flow was measured for 20 min by pilocarpine (0.5 mg/kg) stimulation subcutaneously.

### RNA Sequencing

The raw Quality control of raw paired-end reads sequenced using Illumina platform was performed using FastQC(v0.11.5). Low-quality reads, sequencing adapters, and overrepresented K-mers were removed using Trimmomatic (v0.32), resulting in a total of 694,557,957 trimmed reads (29,978,281 to 120,473,553). The trimmed reads were aligned using the STAR aligner (v2.5.2a) with default parameters to the *Mus musculus* reference genome (Ensemblrelease Mus_musculus.GRCm38.95) to produce BAM files. Raw read counts per gene was generated using HTSeq-Count(v0.6.1p1) with default parameters using GRCm38.95 GTF and library as stranded. The counts generated were filtered to remove unexpressed genes or genes expressed at very low levels to retain only genes with a minimum of five reads in 50% of samples in at least one of the four conditions of interest. Differential gene expression analysis was then performed on the filtered count data (18,594 genes) using R library DESeq2 (1.40.2). Both unsupervised (principal components analysis and correlation analysis) and supervised analyses were done using DESeq2. To calculate transcript abundances, we used Cufflinks v2.2.1 (44) to convert raw reads to FPKM values. Principal component analysis and distribution analysis were used to identify outlier samples. Differential expression analysis was performed using Cuffdiff2 (45). Data visualization was done using JMP Genomics v8 (SAS Institute) and R packages. Two-way hierarchical clustering was done using the Ward method implemented in JMP Genomics. Differences in gene expression were considered statistically significant if the adjusted P value (false discovery rate) was less than 0.05. Pathway enrichment analysis was performed using IPA (Qiagen) software algorithms considering genes differentially expressed between WT and *Stim1/2^K14cre^* cells, with adjusted P < 0.01 and absolute fold change > 1.2 (listed in Supplemental Table 2, A and B). Heatmaps of selected genes were created using the conditional formatting tool in Microsoft Excel. Raw and processed data were deposited in GEO (accession # GSE251691).

### Data analyses and statistics

All statistical analyses were done using GraphPad Prism 9 (GraphPad Software Inc.). Results were expressed as mean ± standard error (SEM). For cell-based assays, at least triplicate measurements were made. Tests for normality (Shapiro-Wilk test) were done prior to performing group comparisons using un-paired Student’s *t*-test parametric or non-parametric Mann-Whitney tests. One-or two-way ANOVA and Student’s *t*-test were used for analysis of in vivo experiments (Saliva flow measurements). Where appropriate, ANOVA was followed by the Tukey’s post-hoc test for multiple comparisons. Where appropriate, we used the Bonferroni for multiple corrections to further analyze differences in the means. Differences with *p*-values of < 0.05 were considered significant: * *p* < 0.05, ** *p* < 0.01, *** *p* < 0.001, and **** *p* < 0.0001. For RNASeq differential expression analysis, an FDR (the Benjamini-Hochberg procedure) of 0.05 was used as threshold for statistical significance.

## Supplementary Material

**Supplemental Figure 1.**
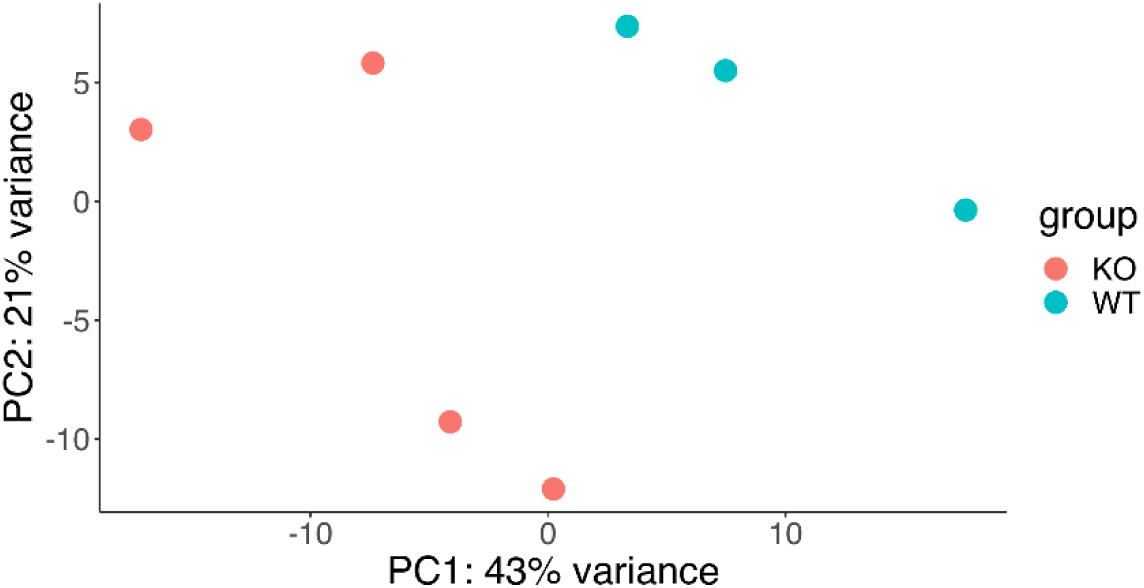
Principal component analysis (PCA) of the normalized RNAseq data obtained from isolated parotid glands in four female mice each of *Stim^1/2K14cre^* (orange dots) and WT (green dots).

**Supplemental Figure 2.**
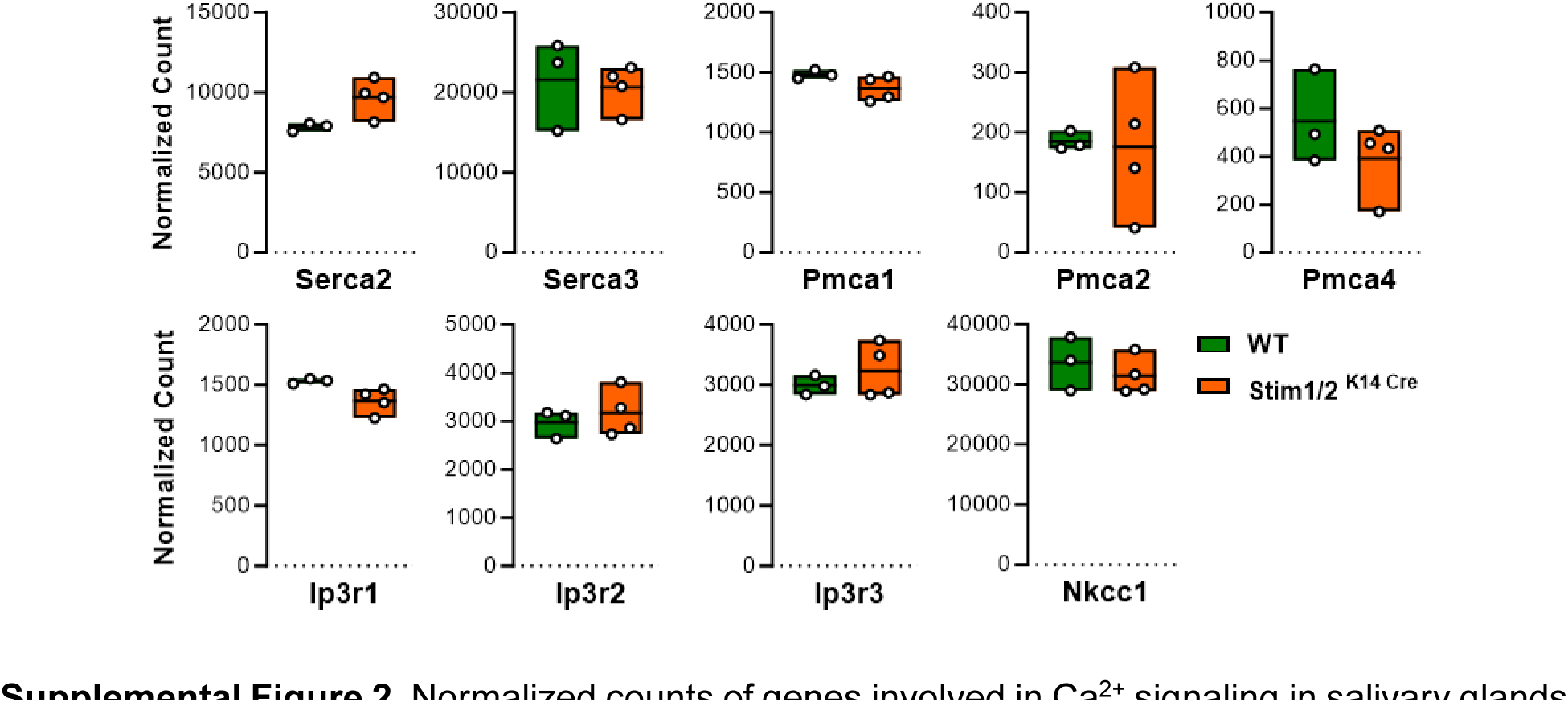
Normalized counts of genes involved in Ca^2+^ signaling in salivary glands such as *Serca2*, *Serca3*, *Pmca1*, *Pmca2*, *Pmca4*, *Ip3r1*, *Ip3r2*, *Ip3r3*, and *Nkcc1* between *Stim1/2^K14cre^* and WT mice. Normalized counts were generated by scaling raw count values to account for sequencing depth using DEseq2 (1.40.2) (*P* < 0.05, One-way ANOVA).

**Supplemental Figure 3.**
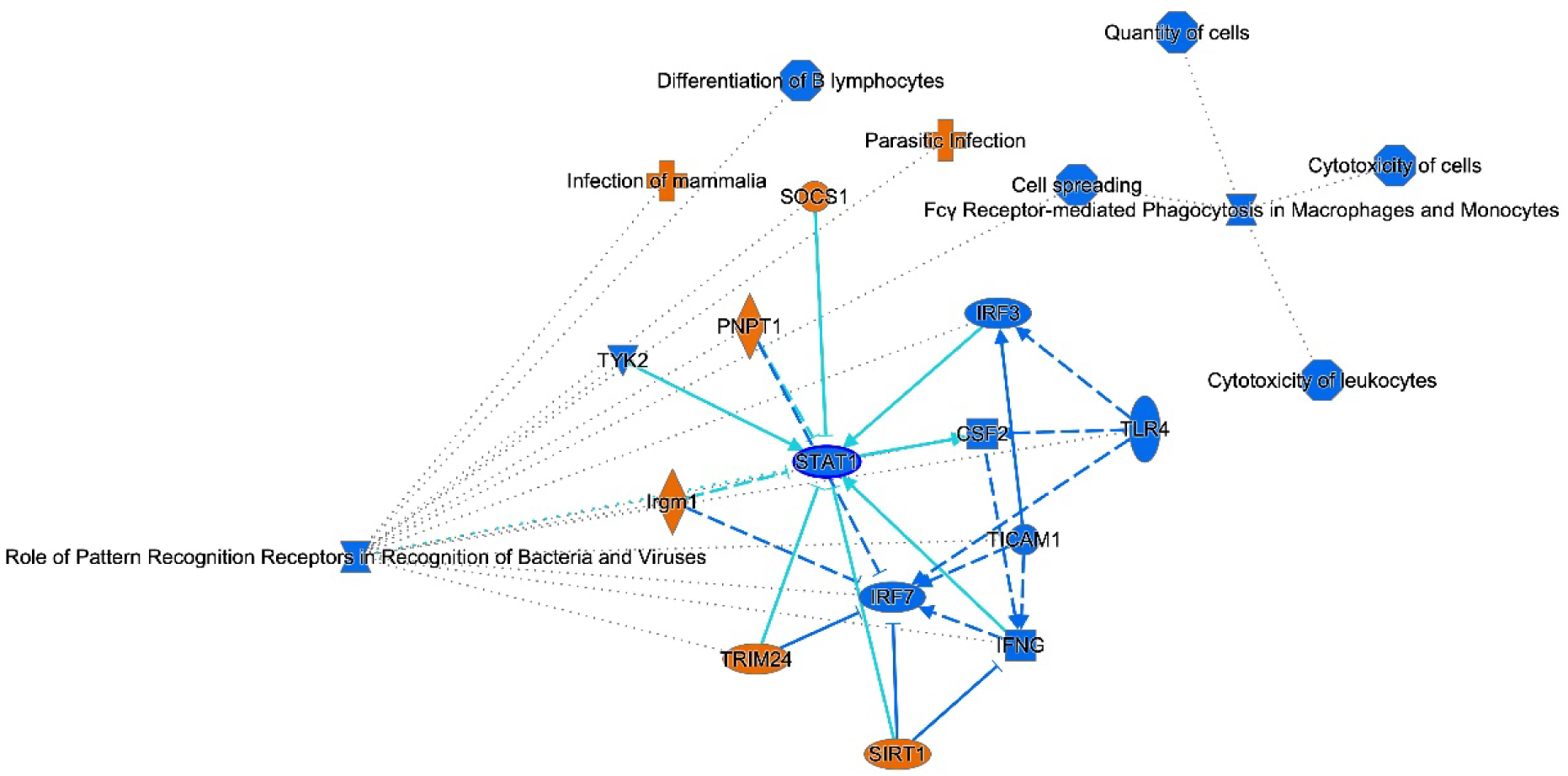
Graphical summary of differentially expressed genes resulting from the comparison of female *Stim1/2^K14cre^* and WT mice. The organizational and functional linkages were derived from Ingenuity Pathway Analysis (IPA) with FDR-CUTOFF ≤ 0.05 and LFC-CUTOFF = ±0.58. Orange indicates predicted activation; blue denotes predicted inhibition.

**Supplemental Figure 4.**
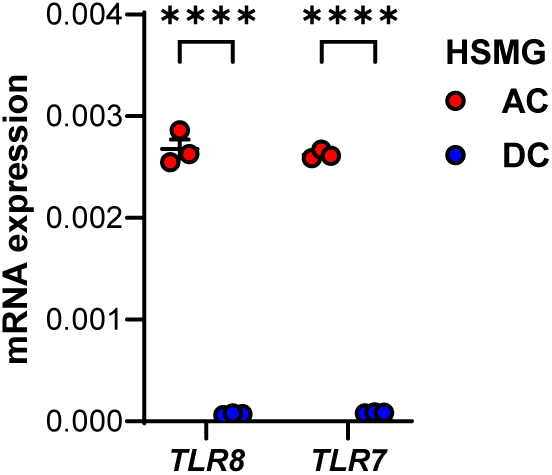
Gene expression analysis of *TLR8* and *TLR7* in the human submandibular acinar cells (HSMG-AC; red dots) and ductal cells (HSMG-DC; blue dots) (2^-ΔCT^). Data represent the mean ±SEM from 3 different experiments (\*\*\*\**P* <0.001, Two-tailed un-paired Student’s t-test).

**Supplemental Figure 5.**
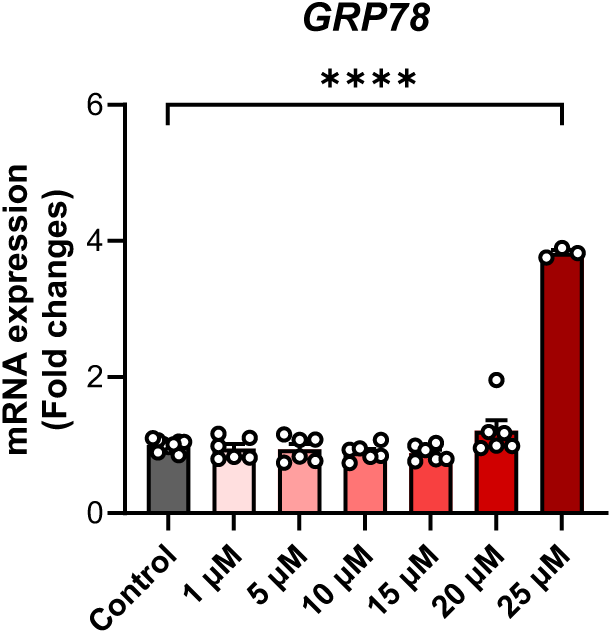
Expression of *GRP78* in CPA treated cells. Human submandibular acinar cells (HSMG-AC) were treated with different concentrations of cyclopiazonic acid (CPA) to reversibly block the sarco-endoplasmic reticulum Ca^2+^ATPase (SERCA). Blocking SERCA can induce endoplasmic reticulum stress. The ER stress maker *GRP78* expression was analyzed at different concentrations of CPA after 2 hrs. Only 25 µM of CPA induced ER stress (\*\*\*\**P* <0.0001, ANOVA).

**Supplemental Figure 6.**
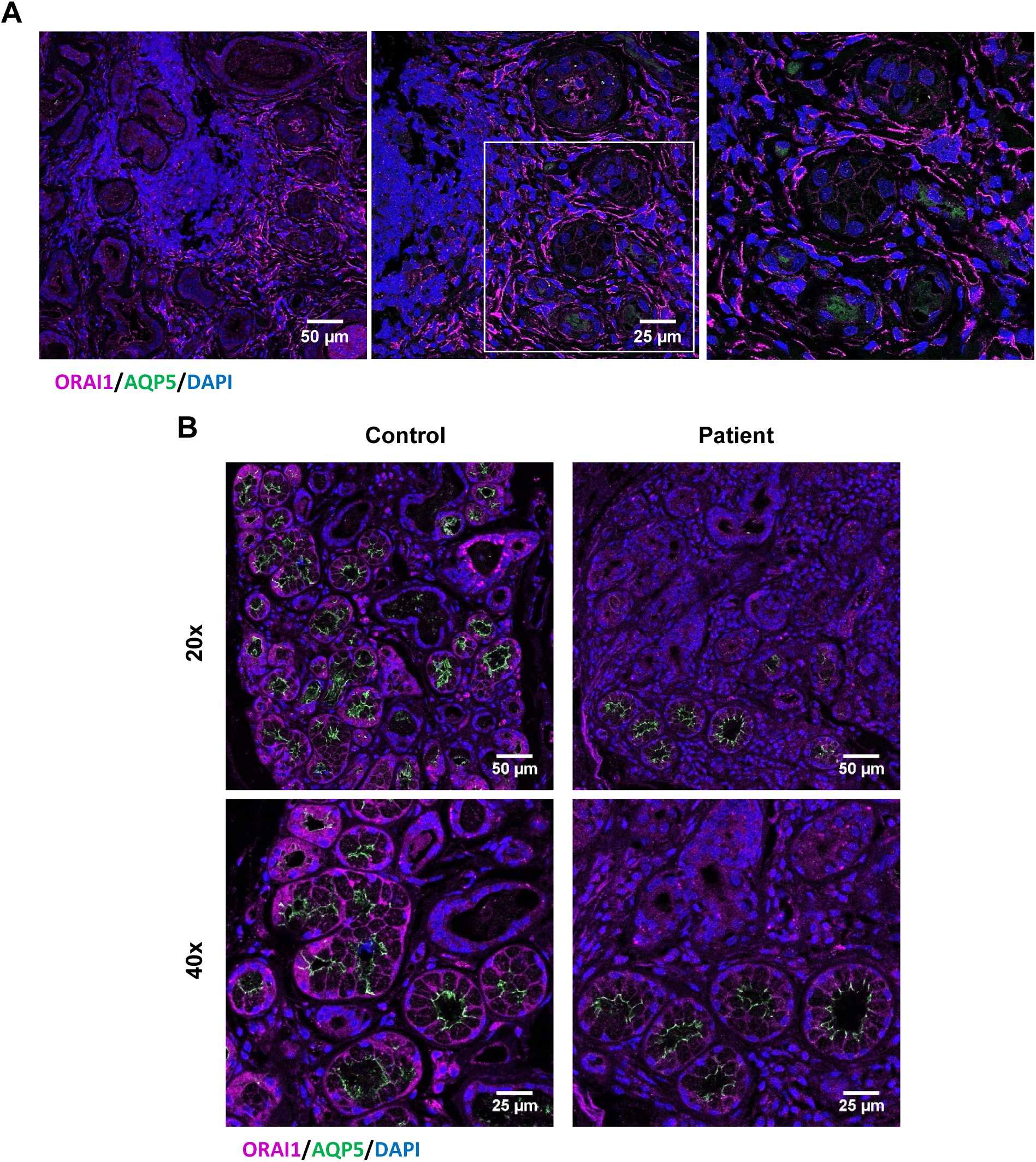
Subcellular localization of ORAI1 and AQP5 in human minor salivary glands from SICCA. **(A)** Representative images of ORAI1 (antibody from the Feske lab), and AQP5 (green) localization in the tissue section of patient minor salivary gland from SICCA. **(B)** Representative images of ORAI1 (purple; Antibody from SIGMA) and AQP5 (green) localization in the tissue section of healthy donor and patient minor salivary gland from SICCA.

**Supplemental Table 1.** Classification of human minor salivary gland samples from SICCA.

| **Samples** | **Score** | **Focus score** | **Baseline Anti-SSA** |
| --- | --- | --- | --- |
| Healthy #1 | 0 | Missing-Skip | Negative |
| Healthy #2 | 0 | Missing-Skip | Negative |
| Healthy #3 | 1 | Missing-Skip | Negative |
| Healthy #4 | 1 | Missing-Skip | Negative |
| Healthy #5 | 1 | 0.4 | Negative |
| Healthy #6 | 1 | 0.4 | Negative |
| Healthy #7 | 1 | 0.5 | Negative |
| Patient #1 | 4 | Missing-Skip | Positive |
| Patient #2 | 4 | 2 | Negative |
| Patient #3 | 4 | 4 | Negative |
| Patient #4 | 5 | 1.2 | Negative |
| Patient #5 | 6 | 0.4 | Positive |
| Patient #6 | 8 | 3.7 | Positive |
| Patient #7 | 9 | 1.4 | Positive |
| Patient #8 | 9 | 2.3 | Positive |

**Supplemental Table 2.**
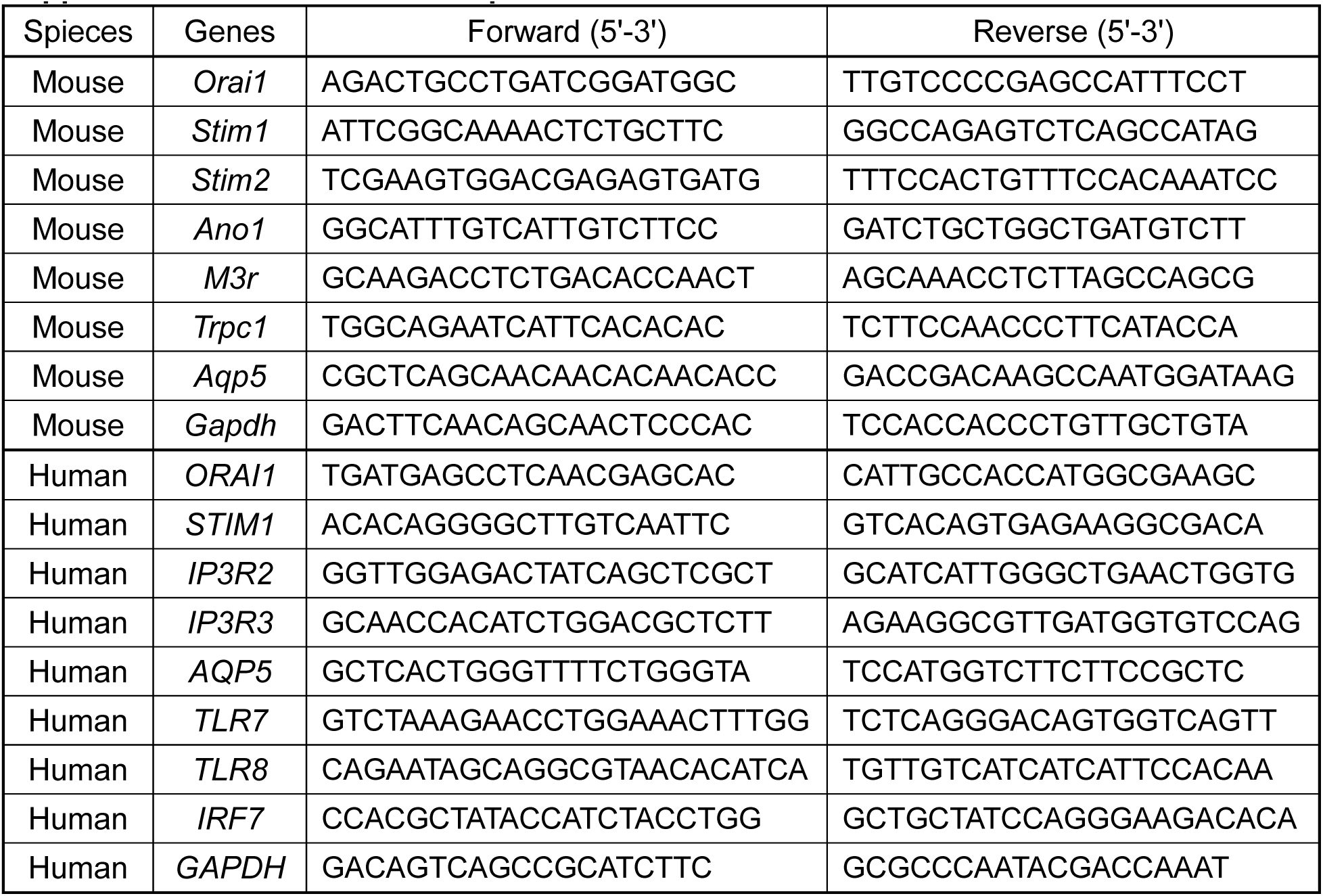
Primer sequences for real-time PCR.

